# Nuclear auxin signaling is essential for organogenesis but not for cell survival in the liverwort *Marchantia polymorpha*

**DOI:** 10.1101/2022.06.21.497043

**Authors:** Hidemasa Suzuki, Hirotaka Kato, Megumi Iwano, Ryuichi Nishihama, Takayuki Kohchi

**Affiliations:** Graduate School of Biostudies, Kyoto University, Kitashirakawa-oiwake-cho, Sakyo-ku, Kyoto 606-8502, Japan; Graduate School of Life Sciences, Tohoku University, 2-1-1, Katahira, Aoba-ku, Sendai, 980- 8577, Japan; Graduate School of Science, Kobe University, 1-1, Rokkodai-cho, Nada-ku, Kobe, 657-8501, Japan; Graduate School of Science and Engineering, Ehime University, 2-5, Bunkyo-cho, Matsuyama, 790-8577, Japan; Department of Applied Biological Science, Faculty of Science and Technology, Tokyo University of Science, Noda 278-8510, Japan

**Author notes:** **Material distribution footnote** The author responsible for distribution of materials integral to the findings presented in this article in accordance with the policy described in the Instructions for Authors (www.plantcell.org) is Takayuki Kohchi.

## Abstract

Auxin plays pleiotropic roles in plant development via gene regulation upon perception by the receptors TRANSPORT INHIBITOR RESPONSE 1/AUXIN SIGNALING F-BOX (TIR1/AFBs). This nuclear auxin signaling (NAS) originated in the common ancestor of land plants. Although complete loss of TIR1/AFBs causes embryonic lethality in *Arabidopsis thaliana*, it is unclear whether the requirement for TIR1-mediated auxin perception in cell viability can be generalized. The model liverwort *Marchanita polymorpha* has a minimal NAS system with only a single TIR1/AFB, MpTIR1. Here we show by genetic, biochemical, and transcriptomic analyses that MpTIR1 functions as an evolutionarily conserved auxin receptor. Null mutants and conditionally knocked-out mutants of Mp*TIR1* were viable but incapable of forming any organs and grew as cell masses. Principal component analysis performed using transcriptomes at various developmental stages indicated that MpTIR1 is involved in the developmental transition from spores to organized thalli, during which apical notches containing stem cells are established. In Mp*tir1* cells, stem-cell- and differentiation-related genes were up- and down-regulated, respectively. Our findings suggest that, in *M. polymorpha*, NAS is dispensable for cell division but essential for three-dimensional body plans by establishing pluripotent stem cells for organogenesis, a derived trait of land plants.

## Introduction

The common ancestors of land plants diverged from algal sisters about 4.5 million years ago and acquired three-dimensional (3D) bodies with parenchymal cells which contributed to their survival in the terrestrial environment (Delwiche and Cooper, 2015). The common ancestor also established several intercellular communication mechanisms mediated by plant hormones (Bowman et al., 2019). Among them, auxin is proposed to act as a morphogen, whose localization and gradient are critical for developmental aspects such as embryonic patterning (Verma et al., 2021; Liao et al., 2015), organ orientation (Galvan-Ampudia et al., 2020; Guan and Jiao, 2020), and gravitropism (Herud-Sikimic et al., 2021; Su et al., 2017) in plants.

Auxin signal is mainly transduced via the nuclear auxin signaling (NAS) pathway, major players of which are: TRANSPORT INHIBITOR RESPONSE 1/AUXIN SIGNALING F-BOX (TIR1/AFB) subunits of Skp1-Cullin-F-box (SCF)-type E3 ubiquitin ligase complex, which act as auxin receptors, AUXIN/INDOLE-3-ACETIC ACID (AUX/IAA) proteins as transcriptional repressors, and AUXIN RESPONSE FACTOR (ARF) proteins as transcription factors. In the NAS pathway, auxin facilitates the interaction between the leucine-rich-repeat (LRR) domain of TIR1/AFBs and domain II (DII) of AUX/IAAs by filling a hydrophobic cavity in the LRR domain, which in turn promotes the ubiquitination of AUX/IAAs (Gray et al., 1999; 2001; Dharmasiri et al., 2005; Kepinski and Leyser, 2005; Tan et al., 2007). In the absence of auxin, AUX/IAAs interact with ARFs and repress transcription by recruiting TOPLESS (TPL) co-repressors to the target loci of ARF (Kim et al., 1997; Ulmasov et al., 1997; 1999; Tiwari et al., 2003). In other words, in the presence of auxin, TIR1/AFBs promote degradation of AUX/IAA with its DII domain acting as a degron, which in turn enables ARFs to exert transcriptional regulation.

Comprehensive phylogenetic analyses of TIR1/AFBs, AUX/IAAs, and ARFs among land plants and green algae have shown that the NAS pathway was established in the common ancestor of land plants through the acquisition of the TIR1/AFB-AUX/IAA co-receptor mechanism in order to regulate transcriptional regulation by ARFs (Bowman et al., 2017; Flores-Sandoval et al., 2018; Mutte et al., 2018). The loss of all six TIR1/AFB homologs in *Arabidopsis thaliana* mutant lines showed disturbed division patterns and delayed cell divisions which led to inhibition of embryogenesis (Prigge et al., 2020). Transmission of *tir1/afb* sextuple mutations through gametes was unaffected, indicating that TIR1/AFBs do not play an essential role in the development of gametophyte, where only a few cell divisions occur (Prigge et al., 2020).

The moss *Physcomitrium* (*Physcomitrella*) *patens* and the liverwort *Marchantia polymorpha* have been studied as models for gametophyte-dominant species (Rensing et al., 2020; Kohchi et al., 2021). Auxin-dependent interaction between TIR1/AFB and AUX/IAA homologs was demonstrated in *P. patens* (Prigge et al., 2010). Disturbed auxin perception suppresses the differentiation of caulonema in *P. patens* (Prigge et al. 2010), whereas in *M. polymorpha* it leads to the formation of undifferentiated cell masses (Kato et al. 2015). In these studies, auxin perception was inhibited either by knockdown of TIR1/AFBs (Prigge et al. 2010) or by the expression of dominant-negative AUX/IAAs with mutations in DII (Kato et al., 2015), where leaky signal transduction cannot be avoided. For this reason, it is difficult to assess the role of auxin in terms of gametophytic cell survival through the studies described above. Instead, knocking out the TIR1/AFBs is expected to shut down NAS altogether. We chose *M. polymorpha* for our study, as it encodes a minimal set of the NAS components, including the sole TIR1/AFB and AUX/IAA homologs, MpTIR1 and MpIAA respectively (Bowman et al., 2017; Flores-Sandoval et al., 2015; Kato et al., 2015).

Germinated *M. polymorpha* spores divide vigorously to produce unorganized tissues, sporelings, and once single-celled apical stem cells are established, they initiate 3D morphogenesis into thalli which differentiate specialized organs such as the gemma cup (Kohchi et al., 2021; Shimamura 2016). Within the dorsal organ gemma cup, multicellular asexual reproductive propagules, gemmae, are produced from single initial cells, which involves apical stem cell formation (Kato et al., 2020). Previous studies have shown that auxin acts as a mobile signal in the thalli (Gaal et al., 1982; Solly et al., 2017) and that NAS regulates the establishment of body axis in gemma development (Kato et al., 2017; Kato et al., 2018). Here, to understand necessity and roles of NAS in plant development, we analyzed molecular functions of MpTIR1 as an auxin receptor and performed knockout-study of Mp*TIR1* in the liverwort *M. polymorpha*.

## Results

### Mp*TIR1* positively regulates auxin response

To determine whether MpTIR1 is involved in auxin response, wild-type (WT), Mp*TIR1*- overexpressing plants under the Mp*EF1A* promoter (Figure 1A) as well as Mp*tir1* mutants were treated exogenously with varying concentrations of natural (indole-3-acetic acid (IAA)) and synthetic (1-naphthaleneacetic acid (NAA) and 2,4-dichlorophenoxyacetic acid (2,4-D)) auxins. WT plants showed ectopic rhizoid formation and arrested thallus development in the presence of high levels of auxin (Figure 1B, C; Supplemental Figure 1A–D). Consistent with the assumption that MpTIR1 is an auxin receptor, Mp*TIR1*-overexpressing plants exhibited hypersensitivity to auxins; the responses were observed at lower concentrations (Figure 1B, C; Supplemental Figure 1).

**Figure 1.**
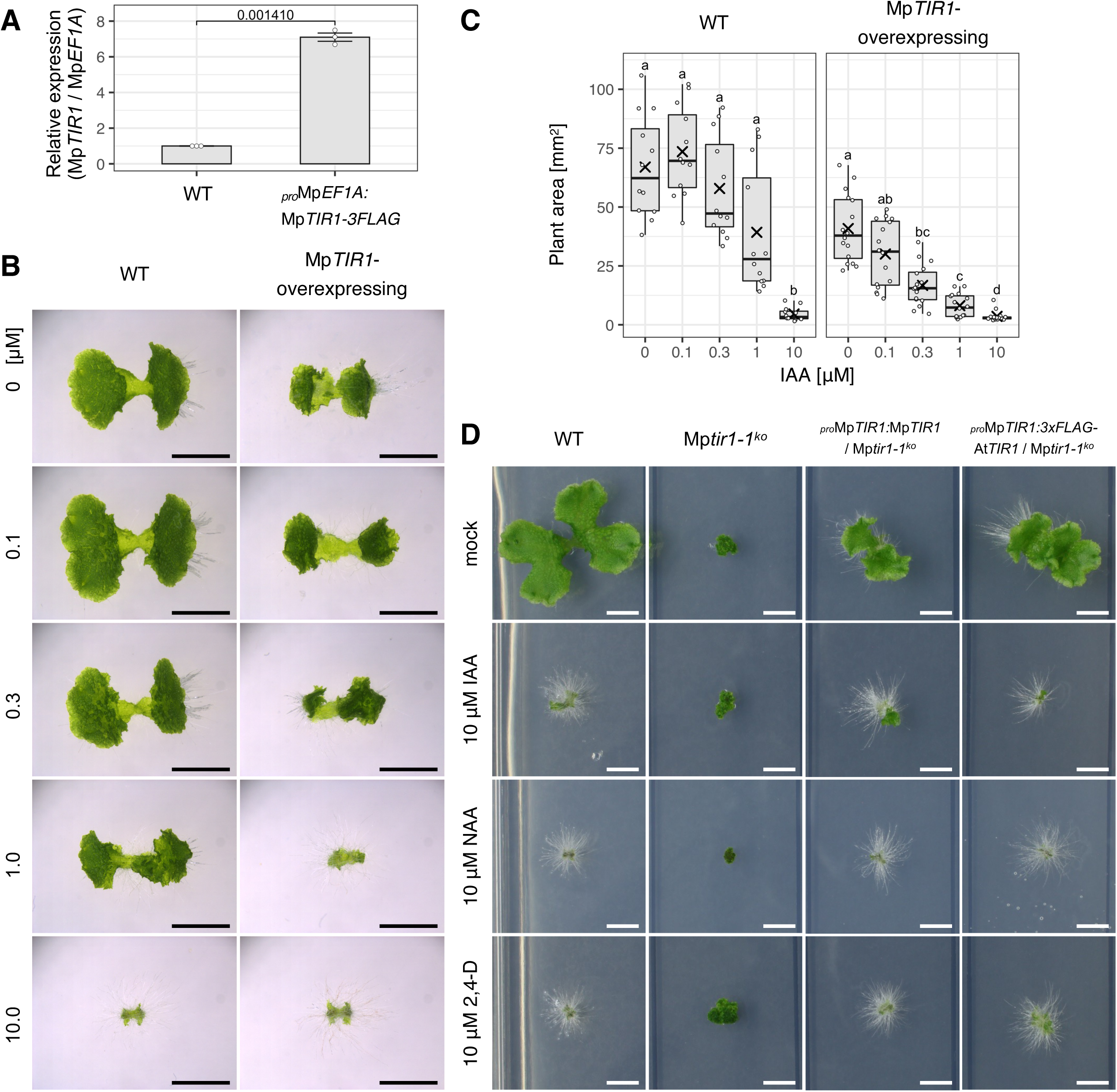
Genetic evidence for Mp*TIR1* as an auxin receptor-encoding gene. **(A)** Relative expression levels of Mp*TIR1* in Mp*TIR1*-overexpressing plants to wild type (WT). Real time-PCR was performed in 10-day-old thalli. Cq values of Mp*TIR1* were normalized by those of Mp*EF1A.* Dots indicate each value of three biological replicates. Error bars indicate mean ± standard error (SE). The value above the plots indicates *p*-value of two-sided Welch’s t-test. **(B, C)** Responsiveness of WT and Mp*TIR1*-overexpressing plants to exogenously supplied auxin. Gemmae were grown on agar media containing the indicated concentrations of IAA for 10 days. (**B**) A representative image is shown for each condition. Scale bars = 5 mm. **(C)** Boxplot of thallus areas. The bands and crosses inside the boxes represent median and mean, respectively. The lower and upper hinges correspond to the first and third quartiles, respectively. Whiskers extend from the hinges to the smallest and the largest values no further than 1.5 * IQR from the hinge (where IQR is the inter-quartile range). Dots represent each value of ≧ 12 biological replicates. Significances were tested by Steel- Dwass test with 99% confidence index. **(D)** Responsiveness of WT, Mp*tir1-1^ko^* mutants, *_pro_*Mp*TIR1:g*Mp*TIR1*/Mp*tir1-1^ko^* plants, and *_pro_*Mp*TIR1:3xFLAG-*At*TIR1*/Mp*tir1-1^ko^* plants to auxin. Small clumps of cell masses (of Mp*tir1-1^ko^*) or gemmae (of the others) were grown for 14 days in the absence or presence of 10 µM each of IAA, NAA, or 2,4-D. Scale bars = 5 mm.

Mp*tir1-1^ko^* mutants grew as cell masses with no obvious organ development (Supplemental Figure2; see “MpTIR1 is critical for organ development but dispensable for cell survival” section for details); exogenous auxins neither caused morphological changes nor enhanced rhizoid development in Mp*tir1-1^ko^* cells, although NAA and 2,4-D, but not IAA, slightly affected the growth increment (Figure 1D; Supplemental Figure 2D, E). Mp*tir1-1^ko^*mutants harboring a transgene of an Mp*TIR1* genomic fragment showed rescued morphology with normal auxin responsiveness, as shown by ectopic rhizoid formation and arrested growth in the presence of high concentrations of auxin (Figure 1D). This was also the case with Mp*tir1-1^ko^* mutants harboring an *A. thaliana TIR1* (At*TIR1*) transgene driven by the Mp*TIR1* promoter (Figure 1D). These results suggest that Mp*TIR1* positively regulates auxin response and that its function is comparable with that of At*TIR1*.

### MpTIR1 acts as an auxin receptor

Interaction between TIR1/AFBs and AUX/IAAs, and the subsequent AUX/IAA degradation are essential for NAS (Gray et al., 1999; 2001; Dharmasiri et al., 2005; Kepinski and Leyser, 2005). To investigate whether MpTIR1 requires auxin for it to interact with MpIAA, we performed an *in vitro* pull-down assay. We used the *Escherichia coli*-expressed fusion protein of glutathione S-transferase (GST) with MpIAA truncated from the 627th amino acid till the C-terminus including DII (GST-MpIAA(627C)) as the bait for *M. polymorpha*-expressed MpTIR1-3xFLAG. The MpTIR1-3xFLAG proteins showed interaction with MpIAA only when auxin was added to the mixture (Figure 2A). IAA facilitated the MpTIR1-MpIAA interaction in a dose-dependent manner (Figure 2A). NAA and 2,4-D also mediated MpTIR1-MpIAA interaction to the same or a slightly weaker degree as IAA (Figure 2A). GST-MpIAA^mutDII^(627C), having a mutated sequence in its DII that had been shown to inhibit auxin signaling in *M. polymorpha* (Kato et al., 2015), did not interact with MpTIR1 regardless of the presence of auxin (Figure 2B). These data suggest that MpTIR1 interacts with the DII of MpIAA in an auxin-dependent manner.

**Figure 2.**
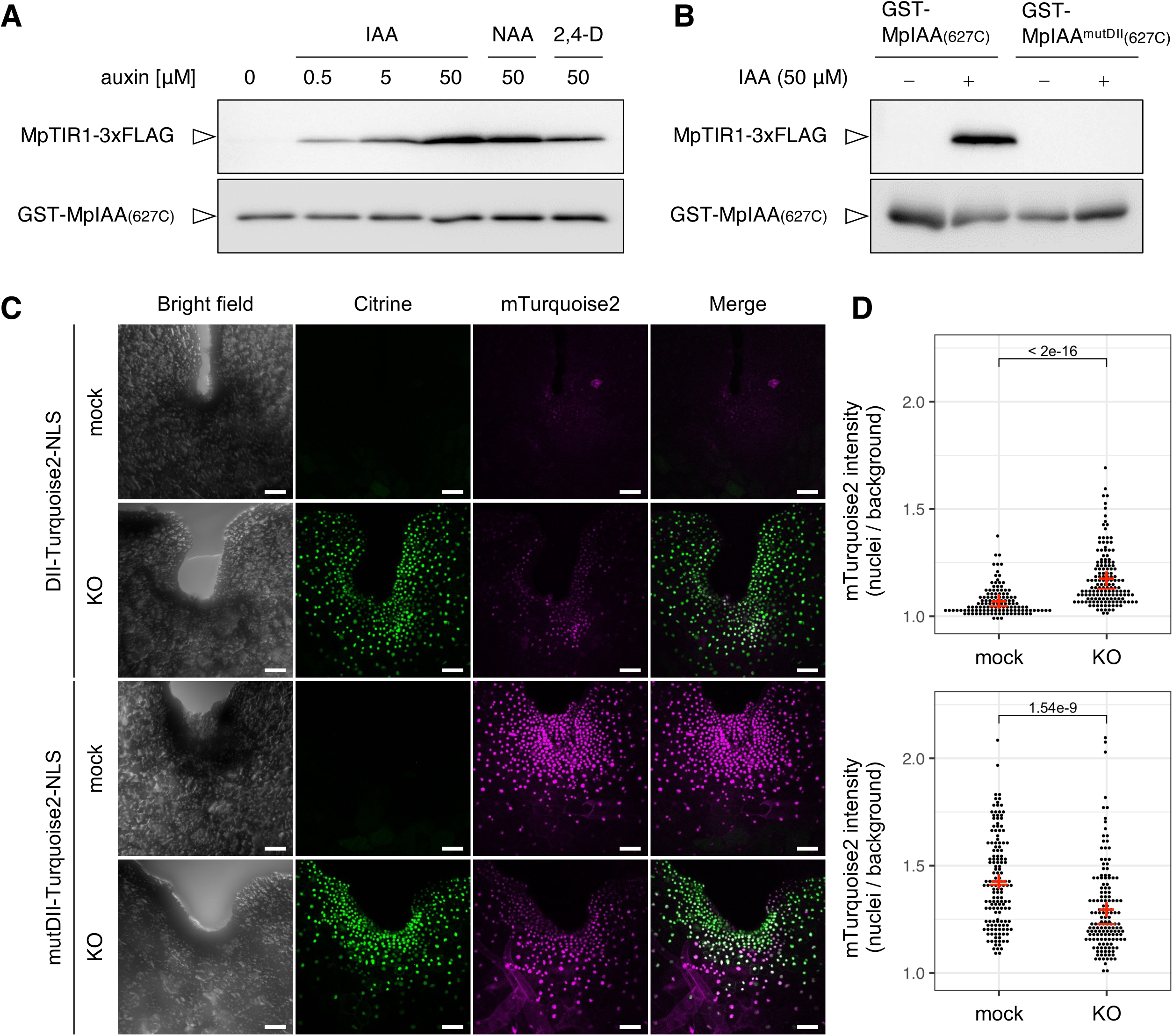
Molecular evidence for MpTIR1 as an auxin receptor. **(A, B)** Pull-down assay between MpIAA and MpTIR1. Bead-bound GST-MpIAA(627-C) proteins, which had been expressed in and purified from *E. coli*, were incubated with a protein extract from MpTIR1-3xFLAG-expressing *M. polymorpha* plants in the presence or absence of the indicated concentrations of IAA, 50 μM NAA, or 50 μM 2,4-D. Bead-bound proteins were washed and subjected to immunoblot analysis with anti-FLAG (top) or anti-GST (bottom) antibody. **(B)** Pull-down assay using a DII-mutated MpIAA. GST-MpIAA(627C) or GST-MpIAA^mutDII^(627C), having a mutation in the DII, were incubated with a protein extract from the MpTIR1-3xFLAG-expressing *M. polymorpha* plants in the presence or absence 50 μM IAA. **(C)** Stabilization of the DII of MpIAA after conditional KO of Mp*TIR1*. One-day-old gemmalings of *_pro_*Mp*EF1A:DII-mTurquoise2-NLS*/Mp*tir1-1^CKO>CitN^* (top) or *_pro_*Mp*EF1A:mutDII- mTurquoise2-NLS*/Mp*tir1-1^CKO>CitN^* (bottom) were either mock-treated or dexamethasone-treated (KO), then subjected to heat shock, and further grown for 1 d. Bright field, Citrine fluorescence, mTurquoise2 fluorescence, and their merged images of a notch region are shown. Scale bars = 50 μm. **(D)** Quantification of mTurquoise2 fluorescence intensities in individual nuclei in the experiments shown in **C**. Dot plots of *_pro_*Mp*EF1A:DII-mTurquoise2- NLS*/Mp*tir1-1^CKO>CitN^* (top) or *_pro_*Mp*EF1A:mutDII-mTurquoise2-NLS*/Mp*tir1-1^CKO>CitN^* (bottom) are shown. The values above the plots indicate *p*-values of Brunner-Munzel test between the mock conditions and KO-induced conditions. The red crosses and bars indicate means and medians, respectively. *n* = 125 or 150 nuclei from 5 or 6 different plants.

We also investigated the involvement of MpTIR1 in the degradation of MpIAA *in vivo* using conditional knockout (CKO) mutants generated by transforming Mp*tir1-1^ko^* cells with a vector (Nishihama et al. 2016) that, under normal conditions, expressed a floxed Mp*TIR1* coding region under Mp*EF1A* promoter for complementation. After induction by heat shock and dexamethasone (DEX) treatment, Cre recombinase excised the floxed Mp*TIR1* to express nuclear localization signal (NLS)-fused Citrine for labeling Mp*TIR1*-KO cells (Supplemental Figure 3A–C). Hereafter, we refer to this as Mp*tir1-1^CKO>CitN^* plants. Then, accumulation of a DII degron peptide probed by mTurquoise2-NLS was assessed in Mp*tir1-1^CKO>CitN^* plants. If MpTIR1 led its target to degradation, accumulation of DII-mTurquoise2-NLS would be facilitated in the absence of MpTIR1. Higher mTurquoise2-fluorescence signals were detected under Mp*TIR1* KO-induced conditions than uninduced conditions (Figure 2C, D). By contrast, the mutated DII peptide (see above) failed to enhance signals of mTurquoise2 fluorescence under Mp*TIR1* KO-induced conditions (Figure 2C, D). These results suggest that MpTIR1 promotes degradation of MpIAA in a DII-dependent manner *in vivo*, which is a direct indication that MpTIR1 acts as an auxin receptor.

### MpTIR1 is essential for auxin-mediated transcriptional regulation

In order to analyze MpTIR1-mediated auxin-dependent transcriptional regulation, WT (5-day-old sporelings) and Mp*tir1-1^ko^*cells were treated with or without 10 μM IAA as well as 10 μM NAA for 4 h and then subjected to RNA-seq analyses to compare transcriptome profiles. Differentially expressed genes (DEGs) were examined in comparison to mock-treated (without auxin) samples for each experiment (Supplemental Data Set 1).

In the WT samples, IAA treatment caused up- and down-regulation of 83 and 76 genes, respectively (Figure 3A). NAA treatment yielded >15-fold larger numbers of DEGs than did IAA treatment (Figure 3A). Nevertheless, the majority of DEGs resulting due to IAA treatment were also differentially expressed in response to NAA with a high correlation (Supplemental Figure 4A, B). These results suggest that although NAA acts more strongly than IAA, both the auxins essentially induce the same transcriptional changes.

**Figure 3.**
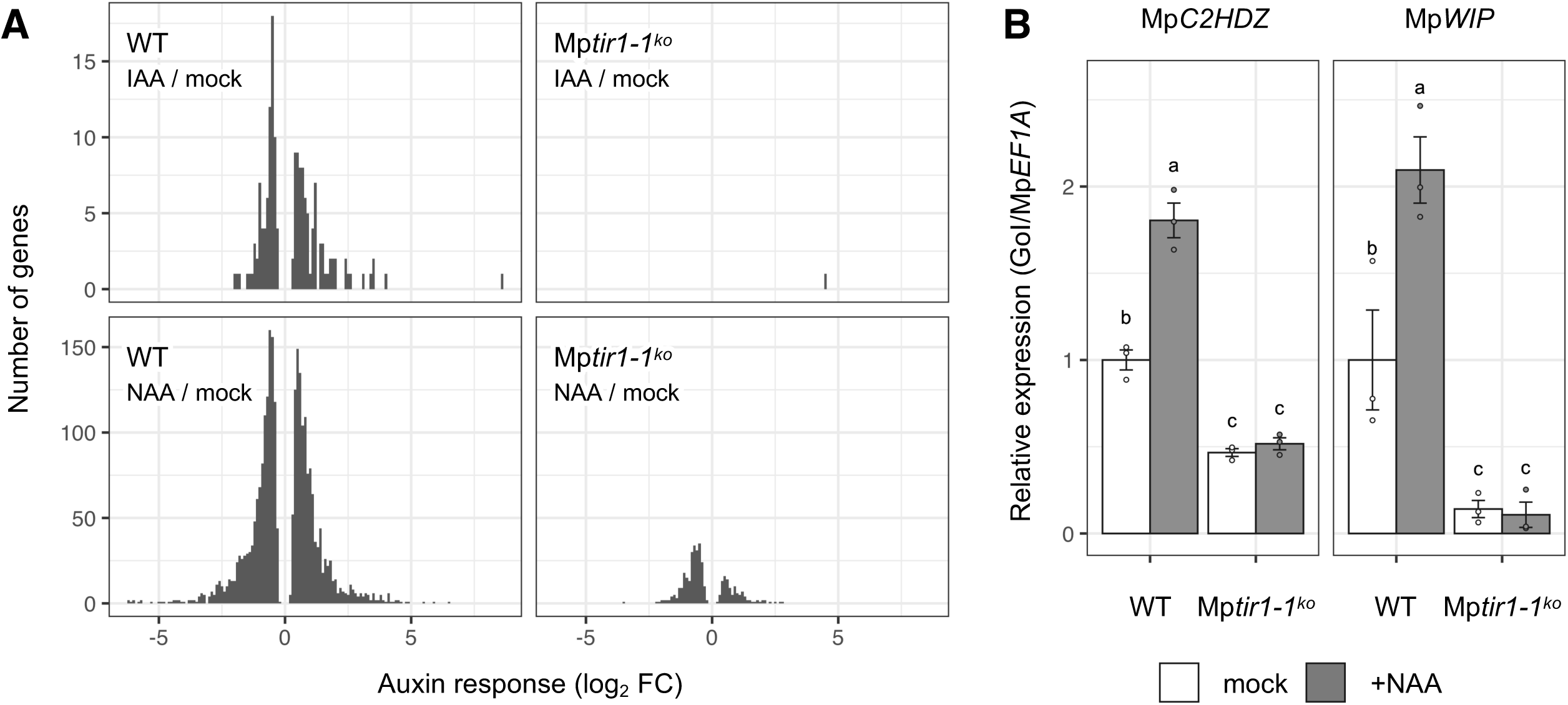
Nearly absolute requirement for Mp*TIR1* in transcriptional responses to auxin. **(A)** RNA-seq analysis of Mp*tir1-1^ko^* cells. X- and y-axes represent log_2_ fold changes (log_2_ FC) and numbers of differentially expressed genes (*p_adj_* < 0.001) in response to IAA (top) or NAA (bottom) treatment in WT sporelings (left) and Mp*tir1-1^ko^* cells (right). Note that the y-axis value of NAA-responsive genes is 10-fold larger than that of IAA. **(B)** Relative expression levels of known auxin-responsive genes, Mp*C2HDZ* and Mp*WIP*, determined by real-time PCR. Five-day-old sporelings and Mp*tir1-1^ko^* cells were treated with 10 μM of NAA or solvent control (mock) for 4 h. Mp*EF1A* was used for normalization. Expression levels relative to mock-treated WT sporelings are plotted. Dots indicate each value of three biological replicates. Error bars indicate mean ± SE. Significances were tested by one-way ANOVA followed by Tukey-Kramer test with 99.9% confidence index.

In Mp*tir1-1^ko^* cells, consistent with MpTIR1 being an auxin receptor, DEGs detected upon IAA treatment were almost nil (up in 1 gene, and down in 0 genes; Figure 3A). DEGs upon NAA treatment (up in 112 genes, and down in 253 genes) were also dramatically fewer than that with WT (Figure 3A). Decreased- or non-responsiveness of Mp*tir1-1^ko^* cells to auxin was further confirmed by real-time PCR on known auxin-responsive genes, Mp*C2HDZ* and Mp*WIP* (Figure 3B; Kato et al., 2017; Kato et al., 2020; Mutte et al., 2018). These results further confirm the role of MpTIR1 in auxin-mediated transcriptional regulation.

### MpTIR1 is critical for organ development but dispensable for cell survival

To understand the role of NAS in 3D morphogenesis, Mp*TIR1* was knocked-out in WT sporelings. The resultant five independent Mp*tir1-1^ko^* mutants failed to develop thalli and slowly proliferated as cell masses (Figure 4A; Supplemental Figures 2A, B, and 5). We could not observe any phenotypic differences between male and female mutants (Supplemental Figure 2C). The mutants displayed unorderly cell division in the cell masses and failed to differentiate into multicellular organs and rhizoids (Figure 4B). To verify these phenotypes, we also genome-edited Mp*TIR1* in WT sporelings (Supplemental Figure 6). Four independent mutants (Mp*tir1- 2^ld^*, Mp*tir1-3^ld^*, Mp*tir1-4^ld^*, and Mp*tir1-5^ld^*), completely lacking the Mp*TIR1* locus, displayed similar phenotypes to those of the Mp*tir1-1^ko^* mutants (Supplemental Figure 7). The developmental defects of Mp*tir1-1^ko^* mutants, besides the auxin response defects described above, were rescued by Mp*TIR1* or At*TIR1* transgenes driven by the Mp*TIR1* promoter (Figure 1D), confirming that the impaired organogenesis of Mp*tir1-1^ko^*mutants was due to loss of evolutionarily conserved functions of MpTIR1. For a more detailed understanding, we compared the cell compositions of Mp*tir1-1^ko^*cell clumps with that of proliferating WT sporelings. Ten-day-old sporelings had meristematic region(s) consisting of small undifferentiated cells and other regions containing large vacuolated cells (Figure 4C, D). Mp*tir1-1^ko^* cell masses were not composed of a uniform cell type but a mixture of small cells at outer regions and large vacuolated cells at inner regions (Figure 4, E, F). Taken together, these results suggest that Mp*TIR1* is dispensable for cell survival but is essential for orderly organogenesis and development.

**Figure 4.**
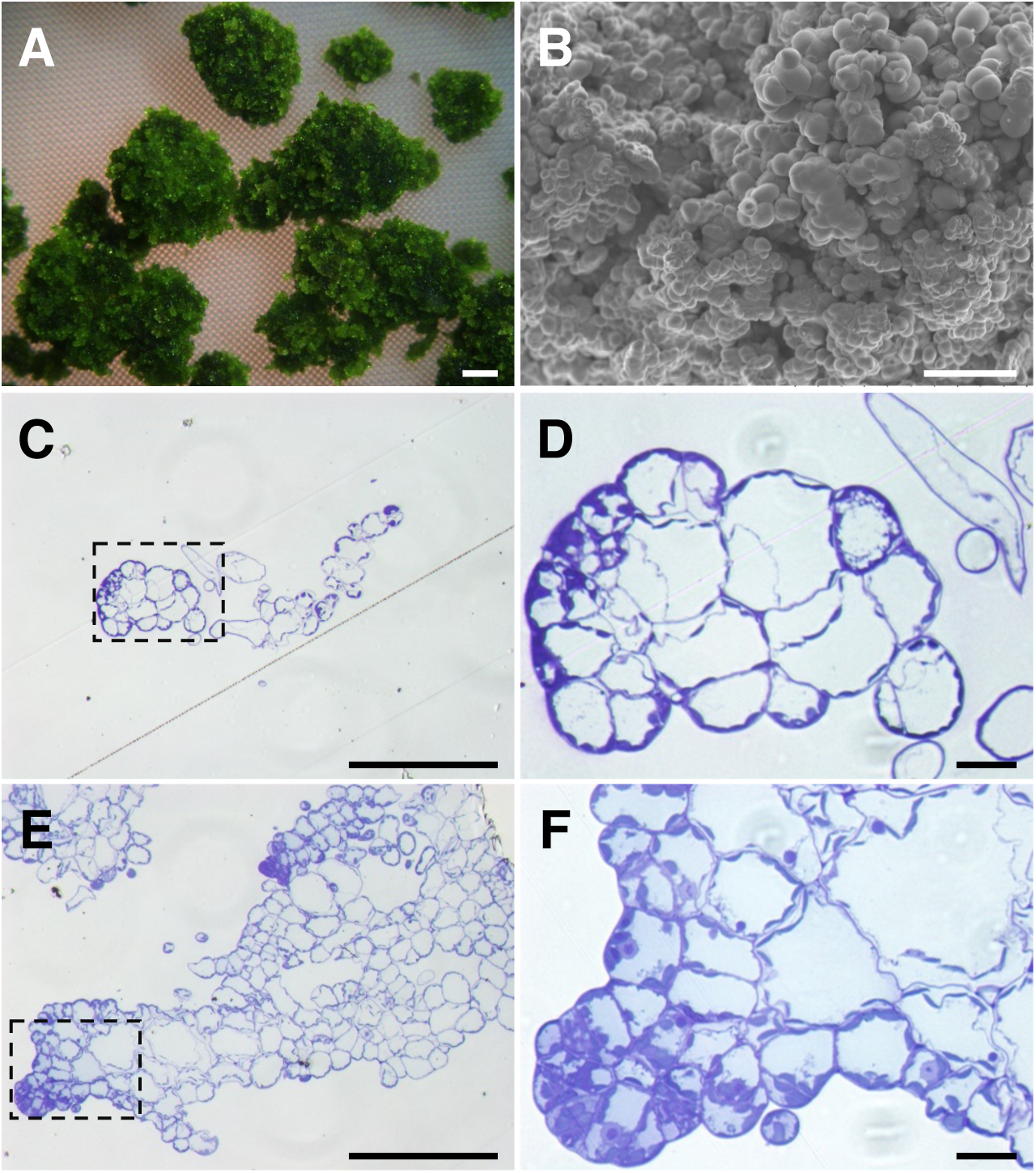
**Requirement for Mp*TIR1* in organogenesis. (A)** Mp*tir1-1^ko^* cell masses. Mp*tir1-1^ko^* cells were grown on an agar medium for 56 days. An overall image is shown in Supplemental Figure S5. **(B)** A SEM image of Mp*tir1-1^ko^* cell masses. A clump of 90-day-old Mp*tir1-1^ko^* cells was observed. (C–F) Section images of WT sporelings (C, D) and Mp*tir1-1^ko^* cell masses **(E, F)**. **(D, F)** Magnified images of the areas within dashed lines in C and E, respectively. Scale bars = 1 mm **(A)**, 200 μm (B, C, E), 20 μm **(D, F)**.

### Mp*TIR1* is critical for proper patterning and organ differentiation

In order to determine the physiological roles of Mp*TIR1* in other developmental stages, Mp*tir1- 1^CKO>CitN^*plants (see “MpTIR1 acts as an auxin receptor” section) and Mp*tir1-1^CKO>tdTN^*plants were generated. In the latter plants, Mp*tir1-1^ko^* cells could be visualized using tdTomato-NLS after excision of the floxed Mp*TIR1* genomic fragment (Figure 5A; Sugano et al. 2018; Suzuki et al., 2020). Mp*tir1^CKO>tdTN^*gemmae essentially showed proper patterning with a trace of stalk at the bottom and two apical notches at its lateral tips (Figure 5B). When we excised the floxed Mp*TIR1* transgene during gemma development, the gemma patterning appeared disordered and failed to establish ellipse-shaped bodies and notches (Figure 5C). We then examined the loss-of- Mp*TIR1* phenotypes after germination of gemmae. Mp*tir1-1^CKO>CitN^* and Mp*tir1-1^CKO>tdTN^* gemmae grew up into thalli under mock conditions, whereas the excision of the floxed Mp*TIR1* transgene in mature gemmae or 1-day-old gemmalings resulted in impaired thallus development and caused cell mass formation (Figure 5D–H, Supplemental Figure 3B, C). These results suggest that Mp*TIR1* is critical for proper patterning and organ differentiation in *M. polymorpha*.

**Figure 5.**
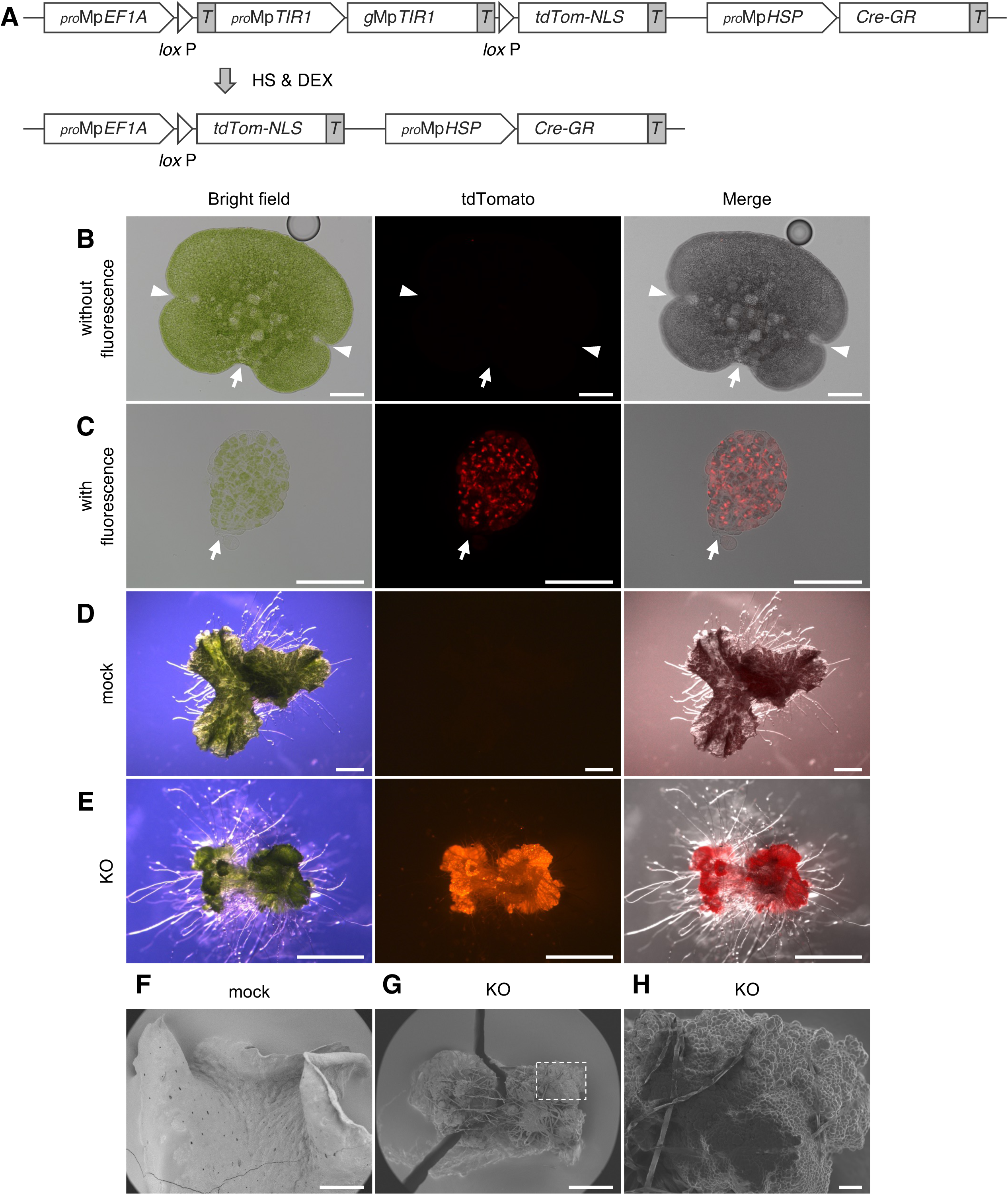
Physiological functions of Mp*TIR1* in establishing the body axis. **(A)** Scheme of the conditional KO (CKO) system in Mp*tir1-1^CKO>tdTN^* plants. An Mp*TIR1* genomic sequence for complementation (top) can be excised by recombination between the flanking *lox*P sequences after heat shock and DEX treatment (bottom), causing the Mp*TIR1* KO situation. *T*: NOS terminator. **(B, C)** Defects in gemma development after CKO of Mp*TIR1*. Mp*tir1-1^CKO>tdTN^* gemmae were grown for 14 days and subjected to KO induction. After further growth for 8 days, gemmae on these plants were observed. Representative gemmae which did not **(B)** or did **(C)** show tdTomato-fluorescence are shown. **(D, E)** Post-germination defects of gemmae after CKO of Mp*TIR1*. Mp*tir1-1^CKO>tdTN^* gemmae were directly subjected to mock-treatment **(D)** or KO-induction **(E)** and further grown for 14 days. **(F-H)** Superficial structures of plants after CKO of Mp*TIR1*. Mp*tir1-1^CKO>tdTN^* gemmae were directly subjected to mock-treatment **(F)** or KO-induction **(G, H)**, further cultured for 22 days, and observed by SEM. **(H)** A magnified image of the area within the dotted line in **G**. Plants with overall tdTomato fluorescence were manually selected as KO-induced samples. Scale bars = 100 μm (**B, C, H**), 2 mm (**D, E**), 1 mm (**F, G**).

### Auxin hypo-responsiveness, a common feature between Mp*tir1-1^ko^*cells and sporelings

In order to clarify the differentiation status of Mp*tir1-1^ko^* cells, we assessed the gene expression patterns using principal component analysis (PCA) of Mp*tir1-1^ko^* cells, sporelings, and available organ-specific transcriptome data (Bowman et al., 2017; Frank and Scanlon, 2015; Higo et al., 2016; Yasui et al., 2019). In the two-dimensional plot of the first and second principal components (PC1 and PC2, respectively), biological replicates from the same sources were clustered (Figure 6A). WT tissues were grouped into an approximate order of developmental stage (Figure 6A). Spores and sporelings were grouped age-wise along PC2, and these developmentally early tissues were grouped separately from the thalli, reproductive organs, and sporophytes along PC1 (Figure 6A). Vegetative thalli and gemma cups were comparable to older sporelings in PC2, while reproductive organs and sporophytes were grouped along PC2 (Figure 6A). Mp*tir1-1^ko^* cells were grouped separately from all of these organs; Mp*tir1-1^ko^* cells were between older sporelings and thalli in PC1 but were similar values to them in PC2 (Figure 6A). These data indicate that Mp*tir1-1^ko^* mutants failed to differentiate from sporelings into mature thalli due to defects in auxin perception.

**Figure 6.**
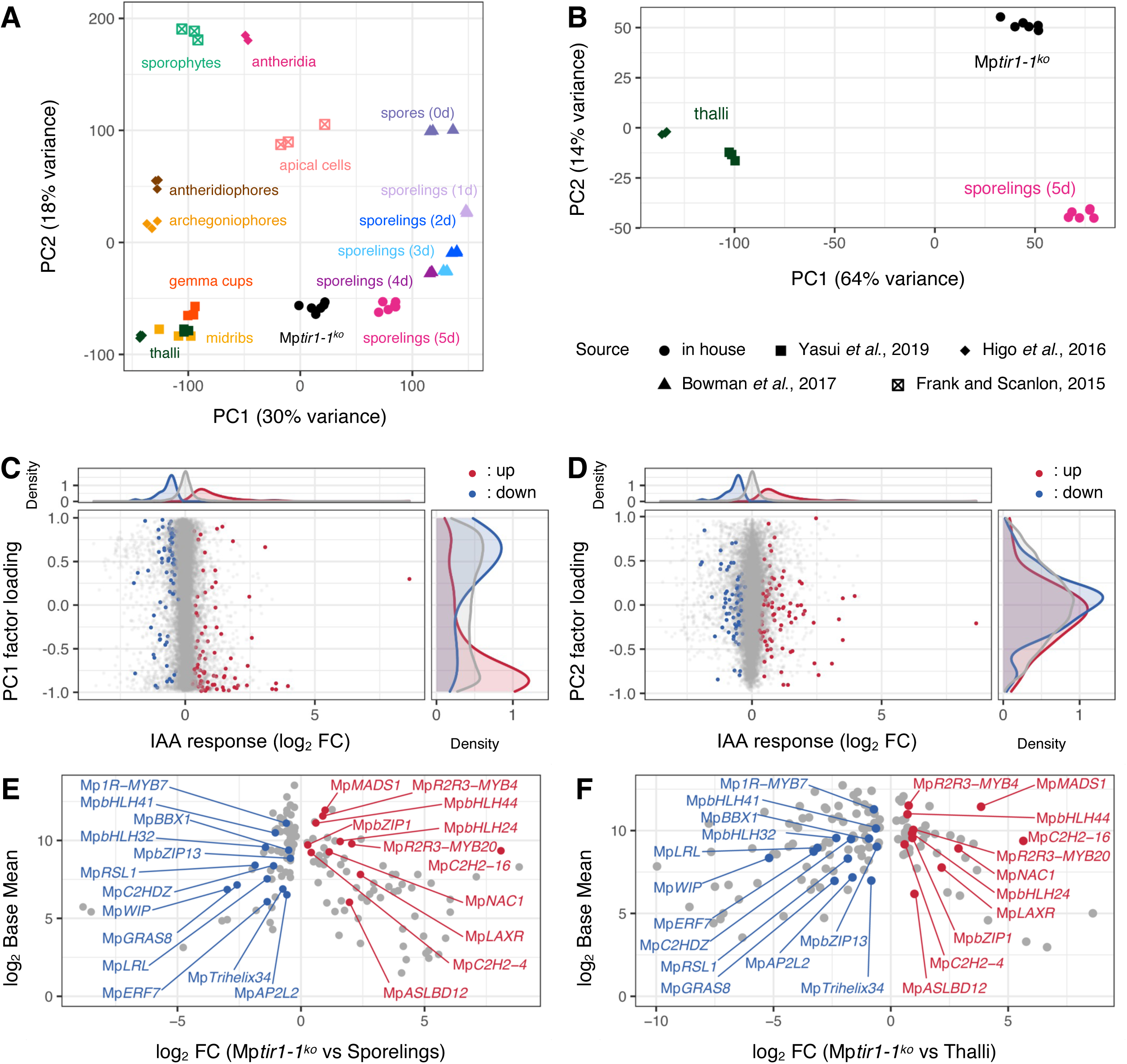
Characteristics of the Mp*tir1^ko^* transcriptome. **(A, B)** PCA of transcriptomes obtained from Mp*tir1-1^ko^* cells and representative *M. polymorpha* tissues. Expression profiles of all genes were used for calculation. **(B)** Re- calculated PCA with a subset including thalli, 5-day-old sporelings, and Mp*tir1-1^ko^* cells. Dots indicate PC1 and PC2 scores of each dataset. Names of tissues or cells are shown near dots in the same color. Dot shapes indicate sources of RNA-seq data. **(C, D)** Contribution of IAA- responsive genes to factor loadings. X-axis of the central panels represents log_2_ FC upon IAA treatment in sporelings. Y-axis of the central panels represents the factor loading of PC1 **(C)** or PC2 **(D)** of the PCA in **B**. Red and blue dots indicate significantly (*p_adj_* < 0.001) up- and down-regulated genes upon IAA treatment, and gray dots represent IAA-non-responsive genes. Top and right panels represent distribution densities of the colored dots along x- and y-axes, respectively. **(E, F)** Expression profiles of transcription factor genes in Mp*tir1-1^ko^* cells. Each dot represents differentially expressed transcription factor genes in Mp*tir1-1^ko^* mutants compared with sporelings **(E)** or thalli **(F)**. Genes found in both comparisons are annotated. X- and y-axes represent log_2_ FC and log_2_ Base Mean (mean of normalized counts of all samples), respectively.

We performed another PCA to confirm this using a subset including Mp*tir1-1^ko^* cells, 5- day-old sporelings, and thalli (Figure 6B). Evidently, PC1 scores were high in Mp*tir1-1^ko^* and sporelings, but low in thalli, resulting in a large separation along PC1 (Figure 6B). In order to find whether auxin contributed to this particular separation, we examined the relationships between auxin responsiveness and factor loadings for PC1 and PC2. Genes that were downregulated upon IAA treatment were dense in positive factor loading of PC1 while those upregulated were dense in the negative factor loading of the PC1 (Figure 6C). These relationships suggest the involvement of IAA in the regulation of gene expression which contributed to high PC1 values in the case of sporelings and Mp*tir1-1^ko^* cells, and low PC1 values in the case of thalli (Figure 6B). In contrast, we did not find any correlation between IAA responsiveness and factor loading of the PC2 (Figure 6D). Similar relationships were observed with respect to NAA treatment (Supplemental Figure 8A, B). These observations suggest that auxin hypo-responsiveness is a characteristic feature of Mp*tir1-1^ko^* cells and sporelings.

### Mp*TIR1* promotes gene expression for organ differentiation

In order to further characterize the differentiation features of Mp*tir1-1^ko^* cells, we performed transcriptomic pairwise comparisons of Mp*tir1-1^ko^* cells with sporelings and thalli (Supplemental Data Set 2). With respect to regulatory genes (Supplemental Data Set 3), Mp*tir1-1^ko^* cells showed significantly lower expression of known transcription factors involved in rhizoid and organ differentiation, such as *LOTUS JAPONICUS ROOTHAIRLESS1-LIKE* homolog (Mp*LRL*; Breuninger et al. 2016), *ROOT HAIR DEFECTIVE SIX-LIKE1* homolog (Mp*RSL*; Proust et al. 2016), and *WIP* homolog (Mp*WIP*; Jones and Dolan, 2017), when compared with that of sporelings and thalli (Figure 6E, F). Conversely, Mp*tir1-1^ko^*cells highly expressed *LOW-AUXIN RESPONSIVE* (Mp*LAXR*), which triggers cellular reprogramming to generate undifferentiated cells (Figure 6E, F; Ishida et al., 2022). Auxin appears to regulate an Mp*TIR1*-dependent expression of these transcription factors (Supplemental Figure 8C–F). Our analyses indicate that Mp*TIR1* plays a critical role in organogenesis by promoting gene expression for differentiation.

## Discussion

### MpTIR1 is an evolutionarily conserved auxin receptor

In this study, we observed a positive correlation between Mp*TIR1* expression levels and auxin responsiveness (Figure 1; Supplemental Figures 1; 2D, E). We observed an auxin-dependent direct interaction of MpTIR1 with DII of MpIAA (Figure 2A, B) which promoted the degradation of DII-tagged proteins (Figure 2C, D). Mp*tir1-1^ko^* mutants were rescued by AtTIR1 (Figure 1D), which further established the role of MpTIR1 as auxin receptor. In *M. polymorpha,* MpTIR1 transmits auxin signal by capturing and leading MpIAA towards degradation, which allows transcriptional activation as well as competitive repression mediated by the sole class-A and class-B ARFs, MpARF1 and MpARF2, respectively (Flores-Sandoval et al. 2015; Kato et al., 2015; Kato et al. 2017; Kato et al. 2020).

Mp*tir1-1^ko^* cells did not show any physiological and transcriptional responses to IAA (Figures 1D; 3; Supplemental Figure 2D, E). This could be due to hyper-accumulation of MpIAA regardless of auxin, which indicates that the transcription of MpARF1/2-target genes are strongly repressed in Mp*tir1* KO cells. Such a constitutively repressive status seems to be different from the absence of A-ARFs, because Mp*arf1* KO mutants showed much milder developmental defects, although the Mp*arf1* mutants were also insensitive to auxin (Kato et al., 2017). The latter situation could be explained by impaired recruitment of MpIAA to MpARF1/2- target loci, as MpARF2 interacts with MpIAA less efficiently than does MpARF1, and thus neither transcriptional activation nor strong repression occurs regardless of auxin (Kato et al., 2017; Kato et al., 2018; Kato et al., 2020). Auxin insensitivity was also caused by loss of all AUX/IAAs as demonstrated in *P. patens* (Lavy et al. 2016), where auxin responses are constitutively saturated in contrast to Mp*tir1* KO cells.

In *M. polymorpha*, the synthetic auxins NAA and 2,4-D inhibit the growth much more severely than does IAA (Ishizaki et al., 2012). In this study, *M. polymorpha* exhibited a much higher number of DEGs with NAA treatment than that with IAA treatment (Figure 3A). However, the amplitude of gene activation or repression by IAA and NAA was comparable with respect to the DEGs regulated by both IAA and NAA (Supplemental Figure 4B). Pull-down assay demonstrated that both IAA and NAA facilitated the interaction between MpTIR1 and MpIAA (Figure 2A). These results suggest that although IAA and NAA both promote MpIAA degradation and enhance transcriptional regulation by MpARF1, we cannot conclude that either of these hormones are functionally stronger. NAA possibly induces more gene expression changes than IAA due to differences in cellular processes such as uptake by passive diffusion (Delbarre et al. 1996) or due to non-specific side effects (Paponov et al. 2019).

### MpTIR1-mediated NAS is essential for 3D morphogenesis but not for cell survival

The observations made in this study where KO mutants of Mp*TIR1* were viable (Figure 4A–C, Supplemental Figures 5, 7), are in line with a previous report indicating *tir1/afb* sextuple mutation did not affect gametophyte viability in *A. thaliana* (Prigge et al., 2020). These results seem to indicate that gametophyte viability is not related to TIR1/AFB-mediated NAS.

In *M. polymorpha*, as is the case for vascular plants, IAA is biosynthesized from tryptophan in a two-step reaction catalyzed first by TRYPTOPHAN AMINOTRANSFERASE OF ARABIDOPSIS (TAA) and then YUCCA homologs (Eklund et al., 2015). Low auxin levels due to KO of the sole TAA homolog MpTAA in *M. polymorpha* or overexpression of IAA conjugation enzyme resulted in cell masses in sporelings (Eklund et al., 2015; Flores-Sandoval et al., 2015). Besides auxin biosynthesis, block of its signaling by dominant suppression of MpARF-mediated gene regulation due to expression of co-repressor-fused MpARFs (Flores-Sandoval et al., 2015) or induction of stabilized MpIAA (Kato et al., 2015) was shown to cause cell mass phenotype. These phenomena are most likely reflected by the cell mass phenotype resulting from Mp*TIR1* KO (Figures 4, 5, Supplemental Figures 3, 7). NAS is proposed to control body axis formation in gemma development as KO of Mp*ARF1* disrupts patternings in this process (Kato et al., 2017). In this study, CKO of Mp*TIR1* in immature gemmae caused structures without notches (Figure 5B, C), probably due to impaired axis formation. These are reminiscent of AtTIR1/AFBs and other downstream elements that control proper patterning in embryogenesis by regulating division orientation in *A. thaliana* (Prigge et al., 2020; Yoshida et al., 2015). These results support that MpTIR1-mediated NAS is essential for 3D body plan establishment and organogenesis during early development.

As cell migration is restricted by cell walls, cell supply from stem cells towards appropriate directions is essential for orderly plant development. The control of cell supply is attributed to the regular division of apical cells in bryophytes (Harrison, 2017; Moody, 2020). In *M. polymorpha*, wedge-shaped apical cells, which produce daughter cells toward the dorsal, ventral, and lateral sides, are established during sporelings and gemma development (Shimamura, 2016).

In the cell mass of Mp*tir1* mutants, establishment of properly shaped apical cells and/or control of division planes may be impaired (Figure 4A, B; Supplemental Figure 7). CKO of Mp*TIR1* in gemmalings where the apical cells are already established also resulted in cell masses (Figure 5D, E, Supplemental Figure 3B, C), suggesting the disruption of apical cell functions in the absence of Mp*TIR1*. PCA based on transcriptomes revealed clear separation of Mp*tir1-1^ko^* cells and sporelings from thalli (Figure 6B) and positive correlation of auxin responsive genes to this separation (Figure 6C, D; Supplemental Figure 8A, B). Mp*tir1-1^ko^* cells showed lower expression of the differentiation-related transcription factors, Mp*LRL*, Mp*RSL,* and Mp*WIP* (Breuninger et al., 2016; Jones and Dolan, 2017; Proust et al., 2016), than sporelings and thalli (Fig6E, F). Judging from this, even though swelled cells were observed (Figure 4F, G), Mp*tir1- 1^ko^* cells were not assumed to be properly differentiated. A relatively mild knockdown of Mp*TAA* results in thalli with impaired organogenesis (Eklund et al., 2015), supporting that auxin response is required for organ differentiation. Mp*tir1-1^ko^* cells showed greater expression of a dedifferentiation-related transcription factor, Mp*LAXR* (Ishida et al., 2022), when compared with that of sporelings and thalli (Figure 6E, F). Although molecular functions have not yet been characterized in *M. polymorpha*, Mp*R2R3-MYB20*, a paralogous gene to *GEMMA CUP-ASSOCIATED MYB1* whose overexpression causes undifferentiated cell clumps (Yasui et al., 2019), and Mp*NAC1*, an orthologous gene to *CUP-SHAPED COTYLEDONs* that act as shoot apical meristem-related boundary genes in angiosperms (Verma et al. 2021), were up-regulated in Mp*tir1-1^ko^* cells. In the moss *P. patens*, auxin signaling is proposed to be low in undifferentiated tissues while it is high in differentiating tissues (Thelander et al., 2019), highlighting the conserved roles of auxin to regulate proper differentiation in gametophyte-dominant species. It could be said that the lack of auxin responsiveness of mutants results in the disruption of apical cell functions, which in turn affects organ differentiation, eventually leading to the formation of undifferentiated cell masses in *M. polymorpha*.

In conclusion, MpTIR1-mediated NAS contributes to establishing 3D body axes through the regulation of apical stem cell functions, including division plane determination and cell differentiation. The findings of this study of Mp*TIR1*, in combination with previous reports of other NAS components in *M. polymorpha*, would help us understand organogenesis through temporal and spatial regulation of NAS in land plants.

## Materials and Methods

### Plant materials and growth conditions

Male accession Takaragaike-1 (Tak-1), female accession Takaragaike-2 (Tak-2), and a female accession of their third backcross generation, BC3-38, were used as wild-type (WT) *M. polymorpha* subsp. *ruderalis*. Tak-1 and BC3-38 were used for phenotypic analysis. Tak-1, BC3- 38, and BC4 spores which were obtained by crossing Tak-1 and BC3-38 were used to generate Mp*TIR1*-overexpressing (*_pro_*Mp*EF1A:*Mp*TIR1-3xFLAG*) plants. F_1_ spores, which were obtained by crossing Tak-1 and Tak-2, were used to generate Mp*tir1-1^ko^* and Mp*tir1^ld^* mutants.

*M. polymorpha* was cultured on half-strength Gamborg’s B5 medium (Gamborg et al., 1968) containing 1% agar under 50–60 μmol photons m^-2^ s^-1^ continuous white light at 22°C unless otherwise defined. For crossing, *M. polymorpha* was grown on soil under far-red irradiated conditions to induce gametangiophore formation as described previously (Chiyoda et al., 2008).

### Preparation of plasmid constructs

Oligos used in this study are listed in Supplemental Table 1.

### For _pro_MpEF1A:MpTIR1-3xFLAG plants

A coding sequence (CDS) of Mp*TIR1* without stop codon was amplified from pENTR_MpTIR1 using the primer pair, MpTIR1_entry/MpTIR1_nonstop, and cloned into pENTR/D-TOPO vector (Thermo Fisher Scientific, Massachusetts, U.S.A.) to generate pENTR_MpTIR1_nonstop. The Mp*TIR1* CDS was then transferred into pMpGWB110 (Ishizaki et al., 2015) by using LR Clonase II (Thermo Fisher Scientific) to generate pMpGWB110_MpTIR1, which was then used for Mp*TIR1*-overexpression experiments and pull-down assays.

### For pull-down assay

Mp*IAA* or Mp*IAA^mutDII^* CDS spanning from the 627th codon till the stop codon was amplified from vectors containing the respective sequences (Kato et al., 2015) using the primer pair, EcoRI-MpIAA_DII/MpIAA-NotI. The PCR products and pGEX6P-1 vector were digested with EcoRI (Takara Bio, Shiga, Japan) and NotI (Takara Bio) and then ligated to generate pGEX6P-1_MpIAA and pGEX6P-1_MpIAAmutDII. Each of these vectors was introduced into *E. coli* Rosetta2(DE3) strain for induction of recombinant proteins.

### For homologous recombination of the MpTIR1 locus

5′- and 3′- homologous arms (3,462 and 3,367 bp, respectively) were amplified from a PAC clone including the Mp*TIR1* locus (pMM23-241G5; Okada et al., 2000) using the primer pairs, MpTIR1_KO_F1/MpTIR1_KO_R1 and MpTIR1_KO_F2/MpTIR1_KO_R2, respectively. The resultant 5′- and 3′- homologous arms were cloned into pJHY-TMp1 (Ishizaki et al., 2013), using the In-Fusion HD cloning kit (Clontech, Mountain View, CA) to generate pJHY- TMp1_MpTIR1, which was then used for homologous recombination of the Mp*TIR1* locus.

### For the complementation cassette for MpTIR1

A genomic fragment spanning from 5,618-bp upstream of the start codon to 1,081-bp downstream of the stop codon was amplified from pMM23-241G5 (Okada et al., 2000) using the primer pair, MpTIR1_usEntry/MpTIR1_R15, and cloned into pENTR/D-TOPO vector (Thermo Fisher Scientific) to generate pENTR_gMpTIR1. The genomic fragment was then transferred into pMpGWB301 (Ishizaki et al., 2015) by using LR Clonase II (Thermo Fisher Scientific) to generate pMpGWB301_gMpTIR1, which was then used for complementation experiments.

### For the expression of AtTIR1 under the MpTIR1 promoter

An Mp*TIR1* promoter sequence spanning from 5,618-bp upstream of the start codon to the start codon was amplified from pMM23-241G5 (Okada et al. 2000) using the primer pair, MpTIR1_usEntry/MpTIR1_R6, and cloned into pENTR/D-TOPO vector (Thermo Fisher Scientific) to generate pENTR_proMpTIR1. An N-terminal 3xFLAG-tagged At*TIR1* CDS was amplified from a vector containing the sequences, pAN19_TIR1 (a sincere gift from Keiko U. Torii and Naoyuki Uchida), using the primer pair, AscI-Flag_F/AtTIR1_AscI_R. pENTR_proMpTIR1 and the *3xFLAG-*At*TIR1* PCR products were digested with AscI (New England Biolabs, Massachusetts, U.S.A.) and then ligated into pENTR_proMpTIR1 to generate pENTR_proMpTIR1:3xFLAG-AtTIR1. The resultant *_pro_*Mp*TIR1:3xFLAG-*At*TIR1* fragment was then transferred into pMpGWB301 (Ishizaki et al., 2015) by using LR Clonase II (Thermo Fisher Scientific) to generate pMpGWB301_proMpTIR1:3xFLAG-AtTIR1, which was then used for expression studies of At*TIR1* under the Mp*TIR1* promoter.

### To generate Mptir1-1^CKO>tdTN^ plants

A *NOS* terminator sequence was amplified from a vector containing the sequence, pMpGWB302 (Ishizaki et al., 2015), using the primer pair, NotI-NosT_F/NotI-NosT_R. The resultant *NOS* terminator fragment and pENTR_gMpTIR1 were digested with NotI (Takara Bio) and then ligated to generate pENTR_NosT-gMpTIR1. The resultant NosT-*g*Mp*TIR1* fragment was then transferred into pMpGWB337tdTN (Sugano et al., 2018) by using LR Clonase II (Thermo Fisher Scientific) to generate pMpGWB337tdTN_NosT:gMpTIR1, which was then used for Mp*TIR1* CKO experiments.

### To generate Mptir1-1^CKO>CitN^ plants

The CDS of Mp*TIR1* was amplified from RNA of WT plants by reverse transcription (RT)-PCR using a primer pair, MpTIR1_entry/MpTIR1_stop, and cloned into pENTR/D-TOPO vector (Thermo Fisher Scientific) to generate pENTR_MpTIR1. The Mp*TIR1* fragment was then transferred into pMpGWB337 (Nishihama et al., 2016) by using LR Clonase II (Thermo Fisher Scientific) to generate pMpGWB337_MpTIR1, which was then used for Mp*TIR1* conditional knockout experiments.

### To generate _pro_MpEF1A:DII-mTurquoise2-NLS/Mptir1-1^CKO>CitN^ plants and *_pro_MpEF1A:mutDII-mTurquoise2-NLS/Mptir1-1^cko>CitN^ plants*

*DII* and *mutDII* sequences of Mp*IAA* were amplified from vectors containing the respective sequences (see Kato et al., 2015) using the primer pair, MpIAA_dN3/MpIAA DII_R1, and then cloned into pENTR/D-TOPO vector (Thermo Fisher Scientific) to generate pENTR_DII and pENTR_mDII. An *mTurquoise2-NLS* fragment was amplified from a vector containing the sequence, pMpGWB337mT2N, using the phosphorylated primer pairs, Aor51HI- mT2_F/NOSt_head_SacI_NLS_mTurq_R, digested with SacI (Takara Bio), and then ligated with Aor51HI- (Takara Bio) and SacI- (Takara Bio) digested pMpGWB203 (Ishizaki et al., 2015) to generate pMpGWB203_Gateway:mT2N. The DII- and mutDII- fragments were transferred into pMpGWB203_Gateway:mT2N by using LR Clonase II (Thermo Fisher Scientific) to generate pMpGWB203-DII-mT2N and pMpGWB203-mutDII-mT2N, respectively. These vectors were then used for degradation assays of DII-tagged protein.

### For locus deletion of MpTIR1 using CRISPR/Cas9 genome editing

Oligos encoding each guide RNA (gRNA) sequence were annealed with their corresponding antisense oligos (see Supplemental Table 1). The annealed oligos were ligated with BsaI (New England Biolabs) digested vectors with the following combinations: 5′gRNA1 and 5′gRNA3 were ligated with pMpGE_En_04, 5′gRNA2 and 5′gRNA4 were ligated with pBC-GE12, 3′gRNA1 and 3′gRNA3 were ligated with pBC-GE23, and 3′gRNA2 and 4′gRNA4 were ligated with pBC-GE34. 5′gRNA1/2-, 5′gRNA3/4-, 3′gRNA1/2-, and 3′gRNA3/4-pairs are active guide RNA-pairs for a nickase version of the CRISPR/Cas9 genome-editing system (Hisanaga et al., 2019; Koide et al. 2020). Four vectors including one of the active 5′gRNA-pairs and one of the active 3′gRNA-pairs were digested by BglI (New England Biolabs) and ligated at once. The resultant gRNA expression cassettes were transferred into pMpGE017 by using LR clonase II (Thermo Fisher Scientific) to generate pMpGE017_MpTIR1_5′gRNA1/2_3′gRNA3/4, pMpGE017_MpTIR1_5′gRNA3/4_3′gRNA1/2, and pMpGE017_MpTIR1_5′gRNA3/4_3′gRNA3/4. These vectors were then used for genome editing of the Mp*TIR1* locus. Mp*tir1-2^ld^* mutation was caused by pMpGE017_MpTIR1_5′gRNA1/2_3′gRNA3/4. Mp*tir1-3^ld^* and Mp*tir1-4^ld^*mutations were caused by pMpGE017_MpTIR1_5′gRNA3/4_3′gRNA1/2. Mp*tir1-5^ld^* mutation was caused by pMpGE017_MpTIR1_5′gRNA3/4_3′gRNA3/4. pMpGE_En04, pBC-GE12, pBC-GE23, pBC-GE34 and pMpGE017 were developed by Keisuke Inoue in Kyoto University.

### Plant transformation

Binary vectors were transformed into WT plants as previously described (Ishizaki et al., 2008 and Kubota et al., 2013). To transform Mp*tir1-1^ko^* mutants, Mp*tir1-1^ko^* cell masses were cocultured with *Agrobacterium* GV2260 harboring binary vectors in liquid 0M51C medium under continuous white light. After 2 or 3 days of co-cultivation, the Mp*tir1-1^ko^* cells were washed with sterilized water and then cultured on half-strength Gamborg’s B5 medium (Gamborg et al., 1968) containing 1% agar, 100 mg/L cefotaxime, and 0.5 μM chlorsulfuron for selection.

### Plant genotyping

For genotyping, small plant fragments were crushed in 100 μL of buffer (100 mM Tris-HCl (pH 9.5), 1 M KCl, and 10 mM EDTA), diluted with 400 µL of sterilized water, and then used as templates for PCR.

Mp*tir1-1^ko^* candidate plants were genotyped by PCR using crude DNA extracts and primer pairs A (MpTIR1_L21/MpTIR1_R12), B (MpTIR1_L14/MpEF_GT_R1), and C (EnSpm_L2/MpTIR1_R13; Supplemental Figure 2A), as described previously (Ishizaki et al., 2013).

Mp*tir1^ld^* candidate plants were genotyped by amplifying the Mp*TIR1* genomic locus from crude DNA extracts using the primer pair, MpTIR1_L30/MpTIR1_R21. The resultant DNA fragments were treated with Exonuclease I and Shrimp Alkaline Phosphatase (New England Biolabs), purified with Fast Gene^TM^ Gel/PCR Extraction Kit (NIPPON Genetics Co., Tokyo, Japan), and then sequenced with the primers, MpTIR1_L45 or MpTIR1_R20.

Molecular determination of sex of the plants was performed by amplifying U- and V-chromosome markers from crude DNA extract, as described previously (Fujisawa et al., 2001) using modified primer pairs, rhf-73F_new/rhf-73R_new and rbm-27F_new/rbm-27R_new, respectively.

### CKO analysis

In order to induce KO of Mp*TIR1* in young plants, 1-day-old Mp*tir1-1^CKO>CitN^*, *_pro_*Mp*EF1A:DII- mTurquoise2-NLS*/Mp*tir1-1^CKO>CitN^*, and *_pro_*Mp*EF1A:mutDII-mTurquoise2-NLS*/Mp*tir1- 1^CKO>CitN^* gemmalings were treated with approximately 3 μL of 10 μM dexamethasone (DEX) solution to enable nuclear localization of glucocorticoid receptor (GR)-fused Cre proteins, dried for several minutes, and then incubated at 37°C for 1 h. Each gemmaling was treated once more with the same procedure at 4-h intervals.

In order to induce KO of Mp*TIR1* in gemmae, Mp*tir1-1^CKO>tdTN^* plant gemmae were treated with approximately 3 μL of 10 μM DEX solution, dried for several minutes, and then incubated at 37°C for 1 h. Each gemma was treated once more with the same procedure at 4-h intervals.

In order to induce KO of Mp*TIR1* in immature gemmae, 14-day-old Mp*tir1-1^CKO>tdTN^* plant thalli were vacuum-infiltrated with 5 μM DEX solution, dried for approximately 20 min, and then incubated at 37°C for 1 h. Each thallus was treated twice with the same procedure at 4- to 5-h intervals. The treated thalli were further cultivated under normal condition for a few weeks. Then, gemmae were taken from gemma cups and used for microscopic analysis.

### Sectioning of plant tissues

In order to observe cell compositions, 10-day-old sporelings and Mp*tir1-1^ko^* cell masses grown on the half-strength Gamborg’s B5 agar media were pre-fixed in 2.5% (v/v) glutaraldehyde and 2% (w/v) paraformaldehyde in 50 mM phosphate buffer (pH 7.2) at 4°C overnight. The samples were then washed with 50 mM phosphate buffer, and treated with 2% (w/v) OsO_4_ solution at room temperature for 2 h, and then washed with 8% (w/v) sucrose. The samples were then dehydrated by 25, 50, and 80% (v/v) ethanol solutions before they were finally dehydrated in 100% (v/v) ethanol, and then embedded into the resin, Quetol 812 (Nissin EM Co., Kyoto, Japan). The resin-embedded samples were sectioned at 1 µm with a diamond knife using Ultracut-UCT (Leica, Wetzlar, Germany). The sections were stained with toluidine blue solution and used for microscopic analyses.

### Microscopic analyses

Thalli, sporelings, and Mp*tir1* cell masses were observed using microscopes SZX16 (Olympus, Tokyo, Japan), M205C (Leica, Wetzlar, Germany), Axiophot (Zeiss, Oberkochen, Germany), or BZ-X710 (Keyence, Osaka, Japan). Plate cultures were photographed using EOS Kiss X3 (Canon, Tokyo, Japan). Z-series images of 2-day-old *_pro_*Mp*EF1A:mutDII-mTurquoise2- NLS*/Mp*tir1-1^CKO>CitN^* plant notches were taken using a confocal microscope FLUOVIEW FV1000 (Olympus). SEM images of Mp*tir1* cell masses and Mp*tir1-1^CKO>tdTN^* plants were taken by TM3000 (Hitachi High Technologies, Tokyo, Japan), as described previously (Nishihama et al. 2015).

### Image manipulations

Bright field (BF) and fluorescence images taken with the M205C were merged using an image analysis software, Fiji (http://fiji.sc/; Schindelin et al., 2012). Z-series confocal images were two-dimensionally projected with max intensity and then merged by using Fiji. BF and fluorescence images taken with BZ-X710 were merged by using BZ-X Analyzer (Keyence).

### Measurement of plant areas

Plant area measurements were performed using Fiji. Bright field images were split into RGB channels by the “Split Channels” function. B-channel images were converted into binary images by the “Auto Threshold” function with the “Default” method. Plant areas were then measured by the “Find Edges” and subsequent “Analyze Particles” functions.

### Quantification of nuclear fluorescence intensities

Images of 2-day-old *_pro_*Mp*EF1A:DII-mTurquoise2-NLS*/Mp*tir1-1^CKO>CitN^* or *_pro_*Mp*EF1A:mutDII- mTurquoise2-NLS*/Mp*tir1-1^CKO>CitN^*plant notches, which were cultured for 1 day after KO induction, were quantified using Fiji by measuring mTurquoise2 fluorescence intensities in the nuclei. Five to six biological replicates were prepared for each condition. A series of 45 confocal images at 5-μm intervals were two-dimensionally projected with sum intensity of the slices. From each image, nucleus and background regions were manually selected as 8-μm circles in diameter at 25 locations each. An average intensity value of each nucleus region was subtracted by mean values of background regions from the same image. The data generated were plotted and was analyzed for statistical significance (see “Statistics and graphics” section).

### Pull-down assay

*E. coli* Rosetta2(DE3) strain harboring the GST-MpIAA(627C) or GST-MpIAA^mutDII^(627C) vectors were precultured in 5 mL of LB liquid medium at 37°C overnight. The overnight grown cultures were then inoculated into 300 mL of fresh LB medium and incubated at 37°C until the OD_600_ reached 0.5. Isopropyl β-D-thiogalactopyranoside (IPTG) was then added to the culture at the final concentration of 0.1 mM for the induction of protein expression; the cultures were incubated at 37°C for 5 h. After 5h, cells were harvested by centrifugation for 10 minutes at 4,000 xg at 4°C, resuspended in ice-cold sonication buffer (PBS and 1 mM dithiothreitol), and subjected to lysis by sonication. The cell lysates were then centrifuged for 30 min at 12,000 xg at 4°C. The supernatants were purified with Pierce^TM^ Disposable Plastic Columns (Thermo Fisher Scientific) of Glutathione Sepharose 4B (GE Healthcare Life Sciences, Massachusetts, U.S.A.).

Fourteen-day-old *_pro_*Mp*EF1A:*Mp*TIR1-3xFLAG* plants were harvested and immediately frozen in liquid nitrogen. The frozen samples were homogenized with three-fourth volume per g of tissue of extraction buffer (150 mM NaCl, 100 mM Tris-HCl pH 7.5, 0.5% Nonidet P-40, 10 μM dithiothreitol, 1 mM phenylmethanesulfonyl fluoride, 1 μg/mL pepstatin A, and 10 μM MG132) and then melted on ice. Debris were removed by centrifugation at 16,000 xg at 4°C for 15 min after which the supernatant was further filtered through a 0.45-μm pore syringe filter. Protein concentration of the samples was measured using Bradford Protein Assay Kit (Bio-Rad Laboratories Inc., California, U.S.A.).

2.5 mg each of the protein samples were incubated with 10 μL of the GST- MpIAA(627C)- or GST-MpIAA(627C)-conjugated Glutathione Sepharose beads and auxin at 4°C for 30 min. After three times of washing with ice-cold extraction buffer, the beads were mixed with 2x Laemmli sample buffer (100 mM Tris-HCl, pH 6.8, 4% [w/v] SDS, 10% [v/v] 2- mercaptoethanol, and 20% [v/v] glycerol), and boiled at 95°C for 5 min. Samples were separated by SDS-PAGE on a 10% acrylamide gel, and transferred onto polyvinylidene fluoride membranes (Bio-Rad Laboratories, Inc.). Membranes were incubated with anti-FLAG (1:5,000; Sigma-Aldrich, Missouri, U.S.A.) or anti-GST (1:2,000; Nacalai tesque, Kyoto, Japan) for 1 h, respectively, washed with PBST (PBS and 0.1% Tween-20), and then incubated with anti-mouse IgG (1:10,000; GE Healthcare) for 1 h. Bands were visualized with ECL Prime reagent (GE Healthcare) and ImageQuant LAS 4010 (GE Healthcare).

### Real-time PCR

For real-time PCR of Mp*TIR1*, 10-day-old Tak-1 and *_pro_*Mp*EF1A:*Mp*TIR1-3xFLAG* plants were harvested and immediately frozen in liquid nitrogen. For real-time PCR of auxin-responsive genes, the F_1_ spores and Mp*tir1-1^ko^* cell masses were precultured in half-strength Gabmorg’s B5 liquid medium for 5 days, treated with 10 μM NAA or solvent control for 4 h, then harvested and immediately frozen in liquid nitrogen. RNA was extracted from the frozen samples using TRIzol (Thermo Fisher Scientific) as described previously (Kubota et al., 2014). Reverse transcription to cDNA and subsequent quantitative PCR were performed as described previously (Kato et al., 2017). Primer pairs, MpTIR1-qPCR_F2/MpTIR1-qPCR_R2, MpC2HDZ-qPCR_F1/MpC2HDZ- qPCR_R1, MpWIP-qPCR_F1/MpWIP-qPCR_R1, and MpEF-qPCR_F/MpEF-qPCR_R were used to quantify Mp*TIR1*, Mp*C2HDZ*, Mp*WIP*, and Mp*EF1A* transcripts, respectively (Supplemental Table1). Mp*EF1A* was used as an internal control. Relative expression levels were calculated by Pfaffl’s method (Pfaffl, 2001).

### RNA-sequencing

The F_1_ spores and Mp*tir1-1^ko^*cell masses were precultured in half-strength Gamborg’s B5 liquid medium for 5 days, and then treated with 10 μM IAA or solvent control for 4 h. Plants were then harvested and immediately frozen in liquid nitrogen. RNA extraction from frozen samples was performed using RNeasy Plant Mini Kit (QIAGEN, Venlo, the Netherland). RNA Libraries were prepared using a NEBNext Ultra II Directional RNA Library Prep Kit for Illumina (New England Biolabs) and sequenced as single end reads using the NextSeq500 platform (Illumina, California, U.S.A.). Total RNA was extracted from the F_1_ spores and Mp*tir1-1^ko^* cell masses treated with 10 μM NAA or solvent control in the same way. Library preparation and subsequent paired-end RNA-sequencing was performed by Macrogen Japan (Tokyo, Japan) using the NovaSeq6000 platform (Illumina).

### RNA-seq data analysis

For quality control, raw read data were pre-filtered using fastp (version 0.20.1; Chen et al., 2018) with default settings for SE- and PE-sequence data, respectively. The filtered reads were then mapped onto the *M. polymorpha* genome (v5.1r1 + U-chromosomal genes of v3.1) using STAR (version 2.6.1c; Dobin et al., 2013) with default settings for SE- and PE-sequence data, respectively. Following analyses were performed in R (version 4.0.0; R Core Team, 2020). Reads mapped on exons were counted using the “featureCounts” function of the Rsubread package (version 2.2.2; Liao et al., 2019). Pairwise comparisons were performed by Wald test using the DESeq2 package (version 1.28.1; Love et al., 2014). In the pairwise comparisons between Mp*tir1-1^ko^* cells and WT tissues, U chromosomal genes were excluded. All the four combinations of comparisons between Mp*tir1-1^ko^*cells (mock samples for IAA or those for NAA) and public thalli data (9-day-old thalli from Higo et al., 2016 or 7-day-old thalli from Yasui et al., 2019) were performed. The shared DEGs among all comparisons were chosen, since all data sets were derived from different experiments, batch effects were not taken into consideration. PCA was performed by the “prcomp” function of the stats package (version 4.0.0; R Core Team, 2020) with log_2_ transformed read counts of all genes. Factor loadings were calculated as √*l* ∗ ℎ_i_/*u*_i_, where *l*, *h_i_*, and *u_i_* represent the eigenvalues of the covariance, the eigenvectors of each gene, and the square root of variance of each gene, respectively.

### Statistics and graphics

Statistical tests were performed by R (version 4.0.0; R Core Team, 2020). The stats package (version 4.0.0; R Core Team, 2020) was used for Welch’s t-test (Figure 1A) and Pearson’s correlation test (Supplemental Figure 4B). The NSM3 package (version 1.15; Schneider et al., 2020) was used for Steel-Dwass test (Figure 1C; Supplemental Figure 1B, D). The lawstat package (version3.4; Gastwirth et al., 2020) was used for Brunner-Munzel test (Figure 2D). The multcomp package (version 1.4.13; Hothorn et al., 2008) was used for ANOVA and subsequent Tukey-Kramer test (Figure 3B), and Dunnett test (Supplemental Figure 2E). The DESeq2 package (version 1.28.1; Love et al., 2014) was used for Wald test (Supplemental Data Set 1, 2). Graphs were drawn by R using the ggplot2 package (version 3.3.1; Wickham, 2016), the ggsignif package (version 0.6.0; Ahlmann-Eltze, 2019), and the UpSetR package (version 1.4.0; Conway et al. 2017; Gehlenborg, 2019).

## Accession Numbers

Sequence data from this article can be found in the GenBank libraries (http://www.ncbi.nlm.nih.gov) or the MarpolBase (https://marchantia.info) under the following accession numbers: At*TIR1* (AT3G62980); MpTIR1 (Mp6g02750 / Mapoly0035s0062); MpIAA (Mp6g05000 / Mapoly0034s0017); MpARF1 (Mp1g12750 / Mapoly0019s0045); MpARF2 (Mp4g11820 / Mapoly0011s0167); MpC2HDZ (Mp2g24200 / Mapoly0069s0069); MpLRL (Mpzg01410 / Mapoly0502s0001); MpWIP (Mp1g09500 / Mapoly0096s0050); MpTAA (Mp5g14320 / Mapoly0032s0124). Other *M. polymorpha* transcription factors used in RNA-seq analysis are listed in Supplemental Data Set 3.

Transcriptome data obtained in this study are stored at DNA Data Bank of Japan Sequence Read Archive (https://www.ddbj.nig.ac.jp/dra) under project number DRA013690. Other public transcriptome data obtained from the Sequence Read Archive (https://www.ncbi.nlm.nih.gov/sra) are listed in Supplemental Table 2.

## Supplemental Data

Supplemental Figure 1. Genetic evidence for Mp*TIR1* involved in auxin response.

Supplemental Figure 2. Genotyping and auxin responses of Mp*tir1-1^ko^* mutants.

Supplemental Figure 3. Verification of induced Mp*TIR1* KO.

Supplemental Figure 4. Significant overlap between IAA- and NAA-responsive genes.

Supplemental Figure 5. Growth of Mp*tir1-1^ko^* mutants.

Supplemental Figure 6. Generation and genotyping of Mp*TIR1-*locus deletion mutants.

Supplemental Figure 7. Reproducibility of Mp*tir1* defects in Mp*tir1^ld^* mutants.

Supplemental Figure 8. Contribution of auxin-responsive genes to transcriptional properties of Mp*tir1-1^ko^*cells.

**Supplemental Table 1.** Oligos used in this study.

**Supplemental Table 2.** Public RNA-seq data used in this study.

**Supplemental Data Set 1.** Pairwise comparisons between auxin- and mock-treated samples in WT or Mp*tir1-1^ko^* cells.

**Supplemental Data Set 2.** Pairwise comparisons between Mp*tir1-1^ko^* cells and WT samples.

**Supplemental Data Set 3.** All *M. polymorpha* transcription factors tested in RNA-seq data analysis.

## Acknowledgements

We thank Keiko U Torii and Naoyuki Uchida for kindly providing the vector, pAN19_TIR1. We thank Keisuke Inoue for kindly providing the vectors, pMpGE_En04, pBC-GE12, pBC-GE23, pBC-GE34 and pMpGE017. We thank Takefumi Kondo and Yukari Sando for sequencing RNA. We would like to thank Editage (www.editage.com) for English language editing.

## Author contributions

H.S., H.K., R.N. and T.K. designed the research and wrote the paper; H.S., H.K., and M.I. performed research and analyzed the data.

## Supplemental Figures and Tables

**Supplemental Figure 1.**
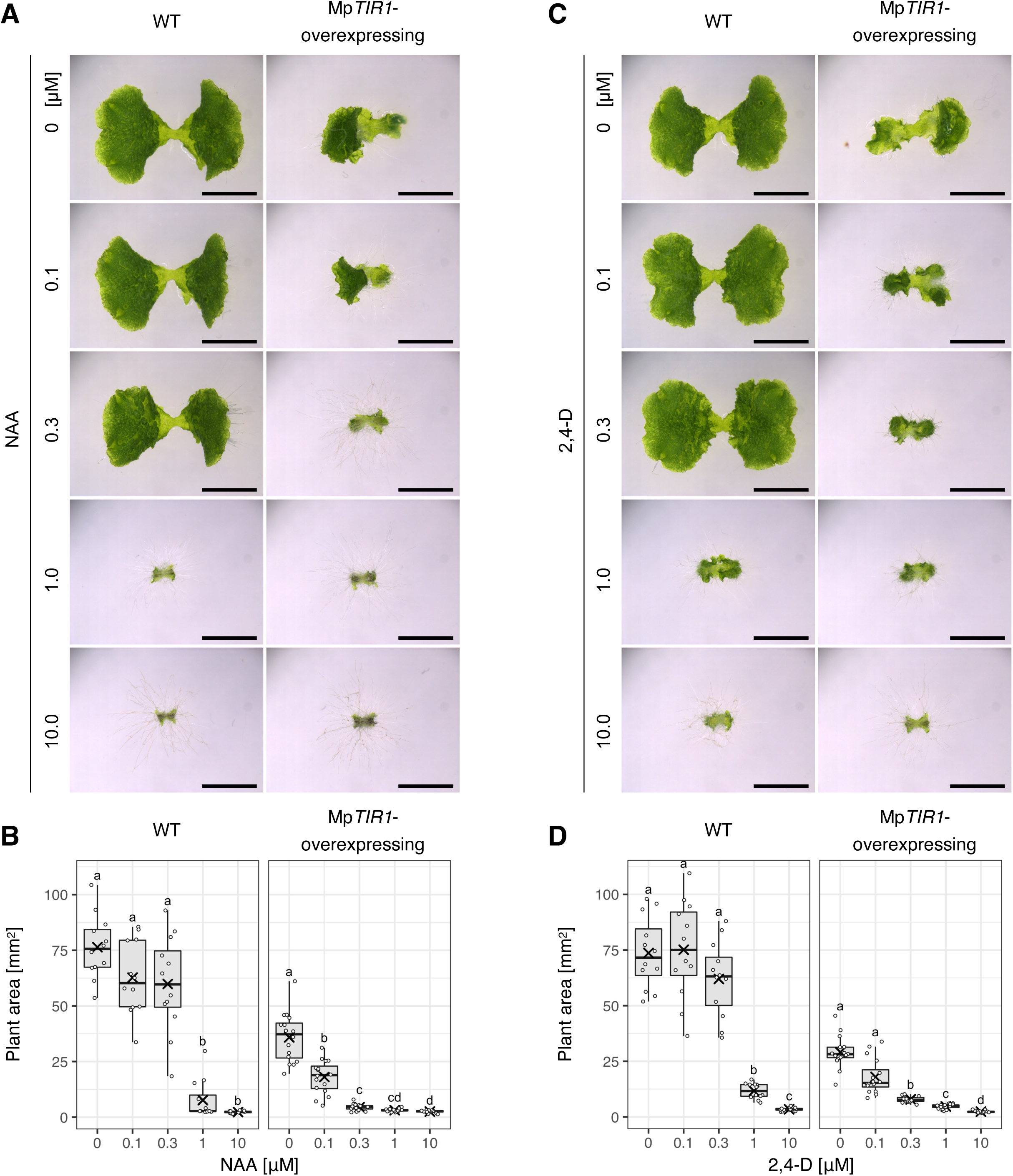
Genetic evidence for Mp*TIR1* being involved in auxin response (Supports. Figure 1**). (A–D)** Responsiveness of WT and *_pro_*Mp*EF1A:*Mp*TIR1-3xFLAG* plants to exogenously supplied auxins. Gemmae were grown on agar media containing different concentrations of NAA **(A, B)** or 2,4-D **(C, D)** for 10 days. (**A, C**) Images of a representative plant for each condition. n ≧ 12. Scale bars = 5 mm. **(B, D)** Boxplot of thallus areas. The bands and crosses inside the boxes represent median and mean, respectively. The lower and upper hinges correspond to the first and third quartiles, respectively. Whiskers extend from the hinges to the smallest and the largest values no further than 1.5 * IQR from the hinge. Dots represent each value of ≧ 12 biological replicates. Significances were tested by Steel-Dwass test with 99% confidence index.

**Supplemental Figure 2.**
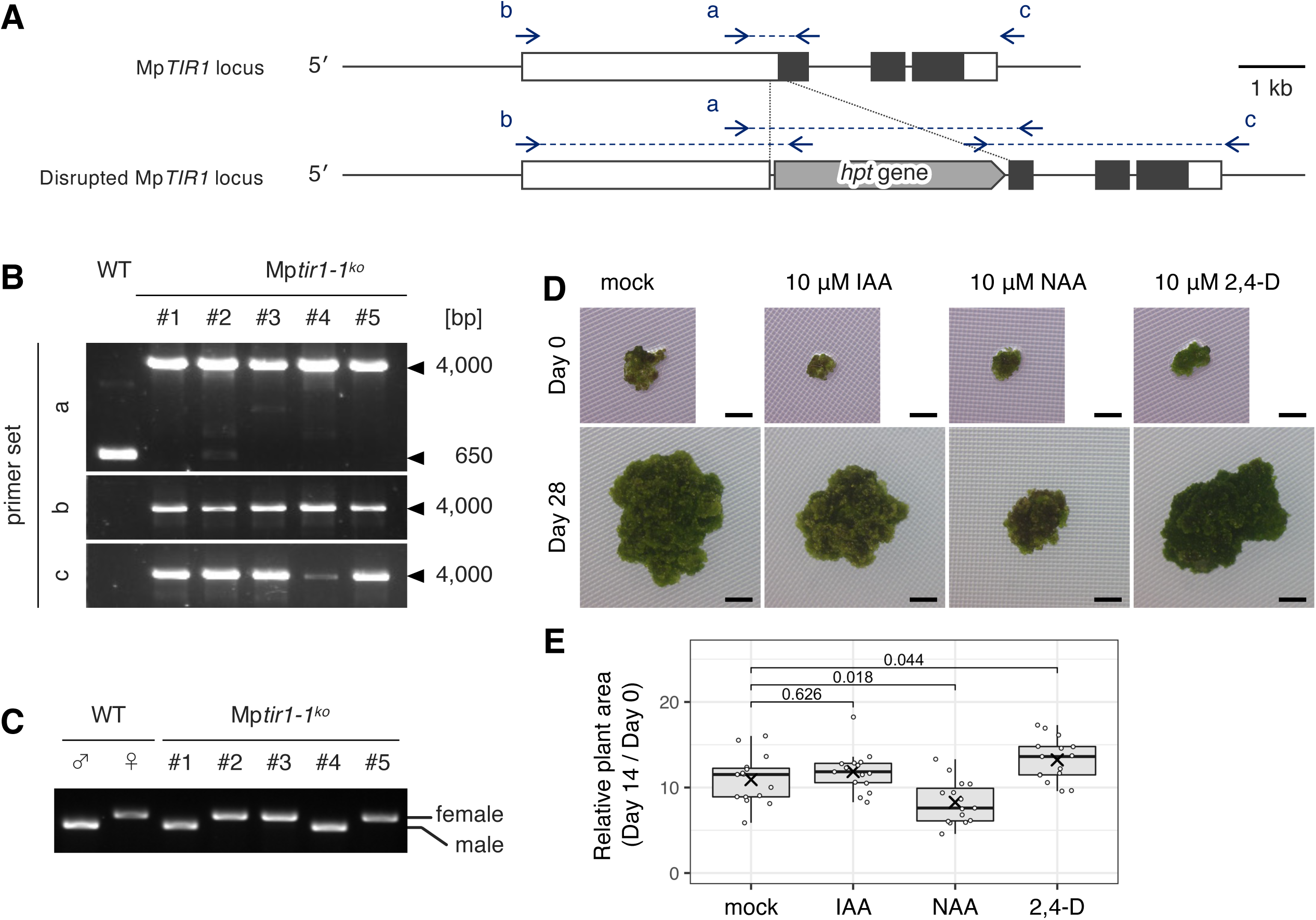
Genotyping and auxin responses of Mp*tir1-1^ko^* mutants (Supports. Figure 1**). (A)** Scheme of the original Mp*TIR1* locus (top) and the homologous recombination disrupted Mp*tir1^ko^* locus (bottom). Arrows with small letters indicate positions of primers or primer sets (connected by a dotted line) used for genotyping PCR. White and black boxes indicate untranslated regions and coding regions, respectively. *hpt* gene: hygromycin phosphotransferase gene cassette. **(B)** Genotyping PCR. The primer sets a–c was used. **(C)** Diagnosis of genetic sex using V (male) and U (female) chromosomal markers. Tak-1 and BC3-38 were used as male and female controls, respectively. **(D, E)** Growth rate of Mp*tir1- 1^ko^* cell clumps in the presence or absence of auxin. Mp*tir1-1^ko^* cell clumps were transplanted on agar media containing 10 μM of various auxins or solvent control (mock) and grown for 14 days. **(D)** Images of a representative clump for each condition. Top and bottom panels show identical plants at 0 and 28 days after transplantation. Scale bars = 1 mm. **(E)** Boxplot of growth rates. Relative expansion of clump areas between day 0 and 14 were calculated as growth rates. The bands and crosses inside the boxes represent median and mean, respectively. The lower and upper hinges correspond to the first and third quartiles, respectively. Whiskers extend from the hinges to the smallest and the largest values no further than 1.5 * IQR from the hinge. Dots represent each value of 15 biological replicates. The values above plots indicate *p*-value of two-sided Dunnett test between mock and auxin-treated samples.

**Supplemental Figure 3.**
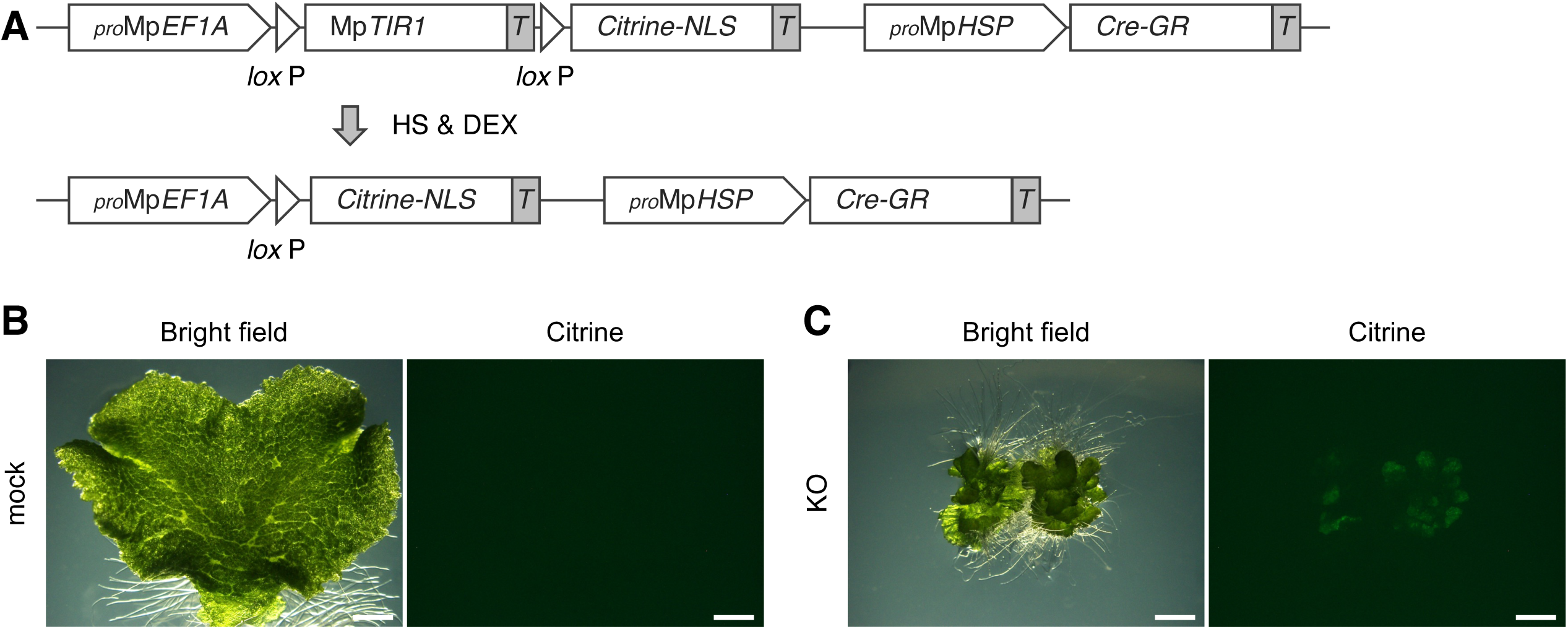
Verification of induced Mp*TIR1* KO (Supports Figure 2 and 6). **(A)** Scheme of the conditional KO (CKO) system in Mp*tir1-1^CKO>CitN^* plants. An Mp*TIR1* genomic sequence for complementation (top) can be excised by recombination between the flanking *lox*P sequences after heat shock and DEX treatment (bottom), causing the Mp*TIR1* KO situation. *T*: NOS terminator. **(B, C)** Phenotype of Mp*tir1-1^CKO>CitN^* plants after KO induction. One-day-old Mp*tir1-1^CKO>CitN^* gemmalings were subjected to mock treatment **(B)** or KO induction **(C)** and grown for 13 days. Scale bars = 2 mm.

**Supplemental Figure 4.**
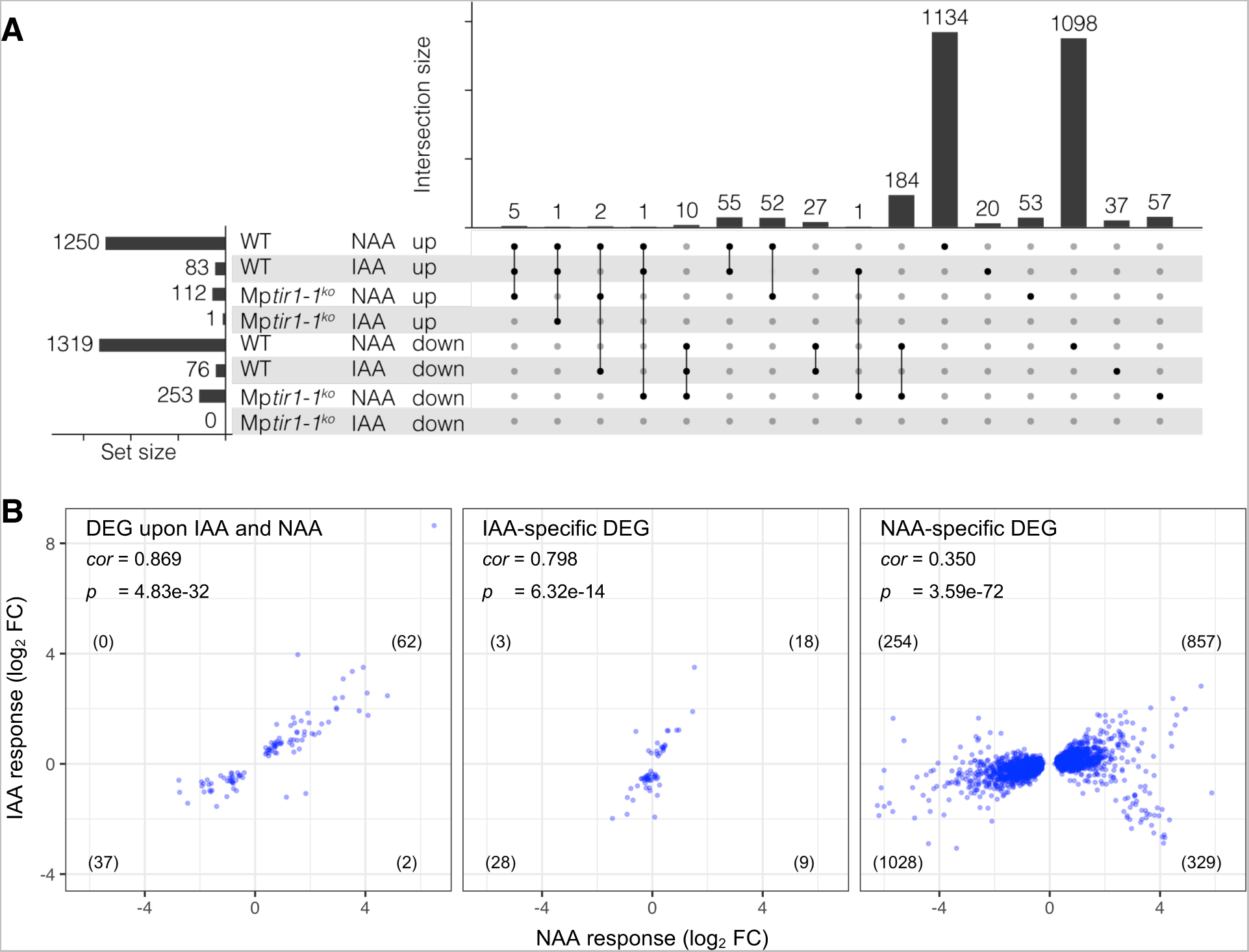
Significant overlap between IAA- and NAA-responsive genes (Supports Figure 3). **(A)** Intersection of auxin-responding genes. Significantly up- or down-regulated genes (*p_adj_* < 0.001) upon IAA or NAA treatment in WT sporelings or Mp*tir1-1^ko^* cells are shown in UpSet plot (Lex et al., 2014). **(B)** Properties of transcriptional responses to different auxins. Log_2_ FC of differentially expressed genes in response to IAA and NAA commonly or specifically in WT sporelings are plotted. *cor* and *p* indicate Pearson’s correlation coefficient and its *p-*value, respectively, between the log_2_ FCs in IAA and NAA responses. The number of genes in each quadrant is shown in parenthesis.

**Supplemental Figure 5.**
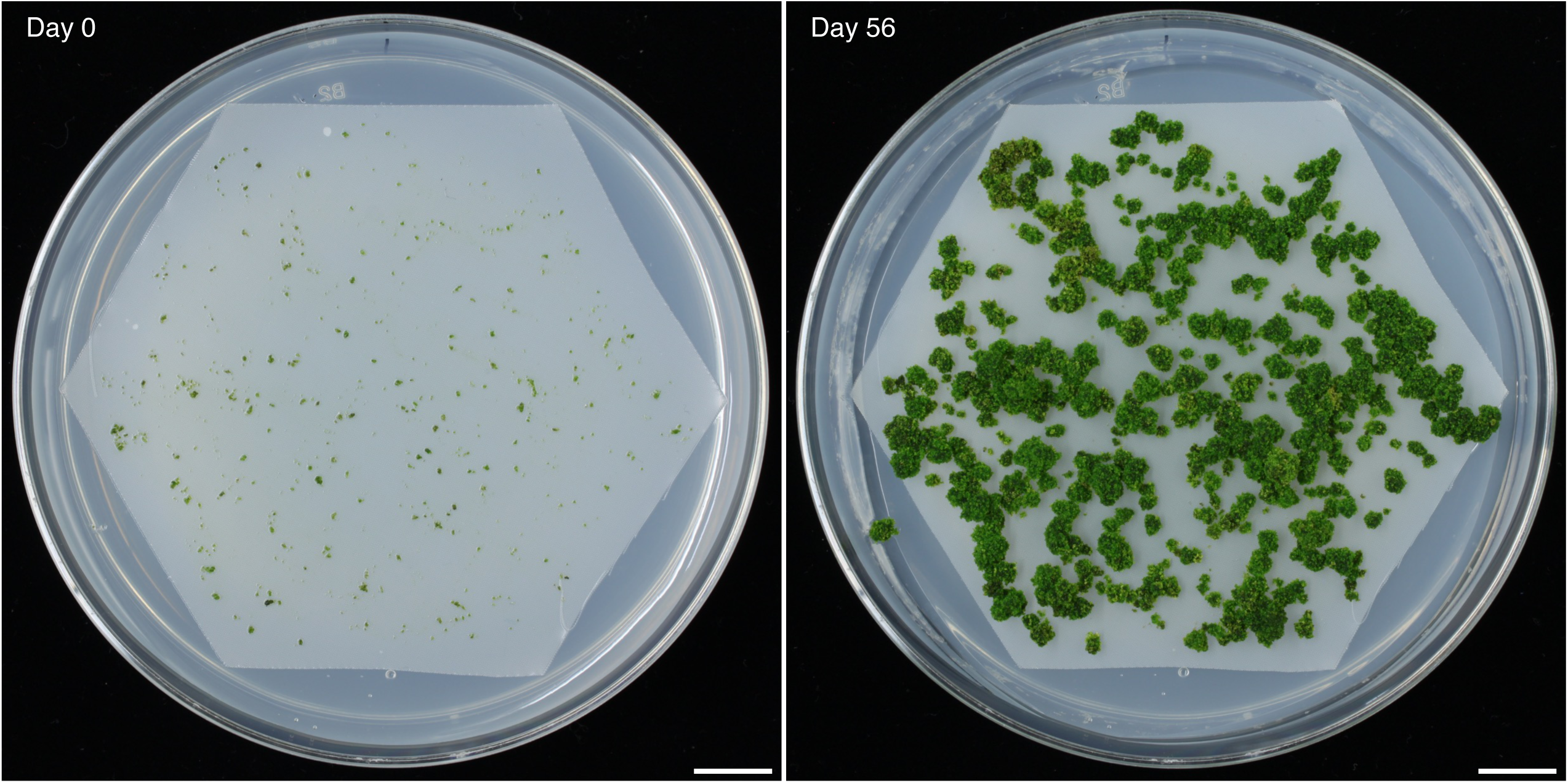
Growth of Mp*tir1-1^ko^* mutants (Supports Figure 4). Mp*tir1-1^ko^* cells were grown on an agar medium covered with nylon mesh. Left and right panels show pictures of an identical plate at day 0 and 56, respectively. Scale bars = 10 mm.

**Supplemental Figure 6.**
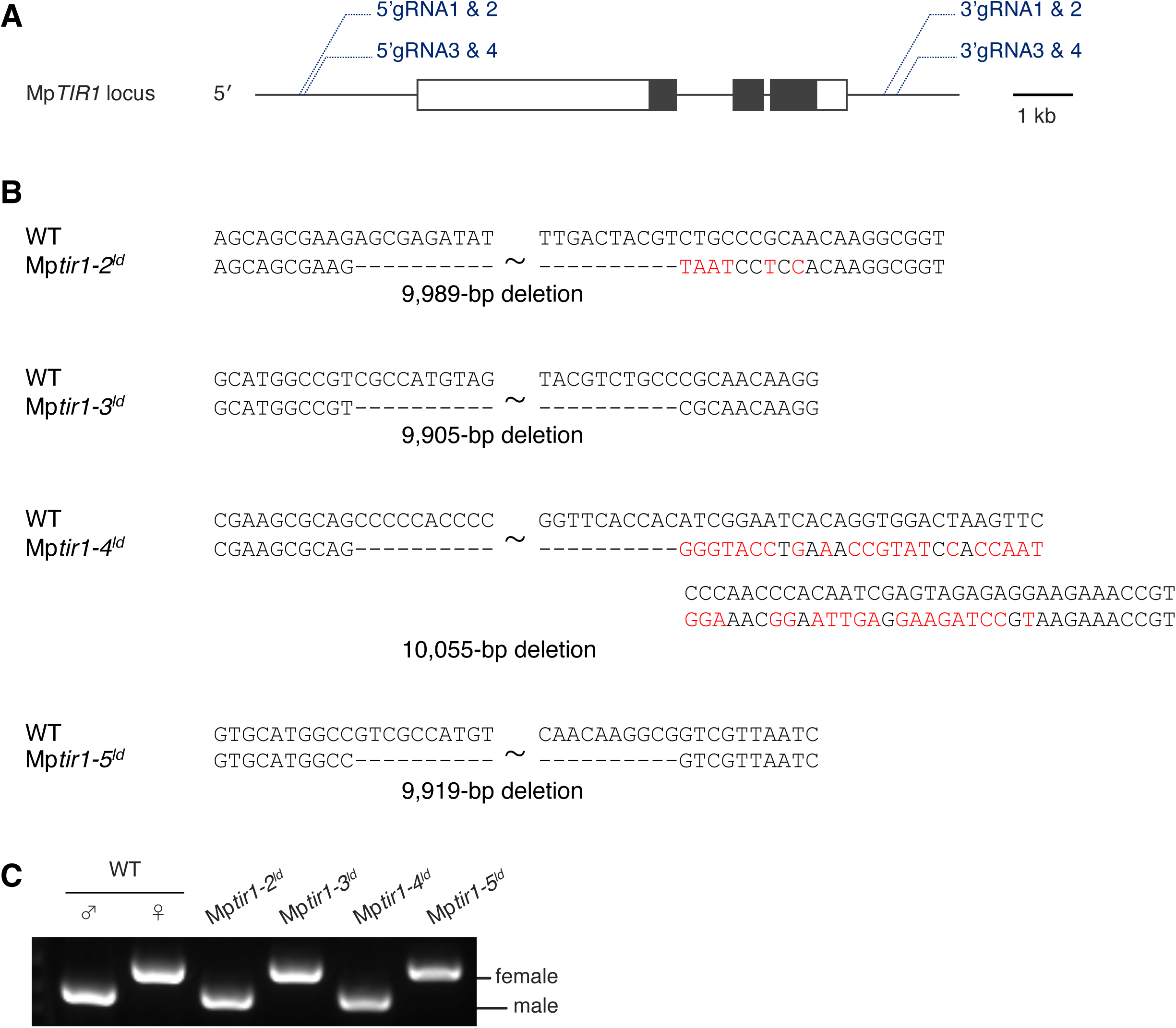
Generation and genotyping of Mp*TIR1-*locus deletion mutants (Supports Figure 4). **(A)** Scheme of the Mp*TIR1* locus and gRNA pairs. 5′gRNA1 & 2 and 3′gRNA3 & 4 pairs were used to generate the Mp*tir1-2^ld^* mutant allele. 5′gRNA3 & 4 and 3′gRNA1 & 2 pairs were used to generate the Mp*tir1-3^ld^* and Mp*tir1-4^ld^* mutant alleles. 5′gRNA3 & 4 and 3′gRNA3 & 4 pairs were used to generate the Mp*tir1-5^ld^* mutant allele. White and black rectangles indicate untranslated regions and coding regions, respectively. **(B)** Mutations in the Mp*TIR1*-locus deletion mutant alleles. Hyphen and red characters indicate deleted and mutated nucleotides, respectively. **(C)** Diagnosis of genetic sex using V (male) and U (female) chromosomal markers. Tak-1 and BC3-38 were used as male and female controls.

**Supplemental Figure 7.**
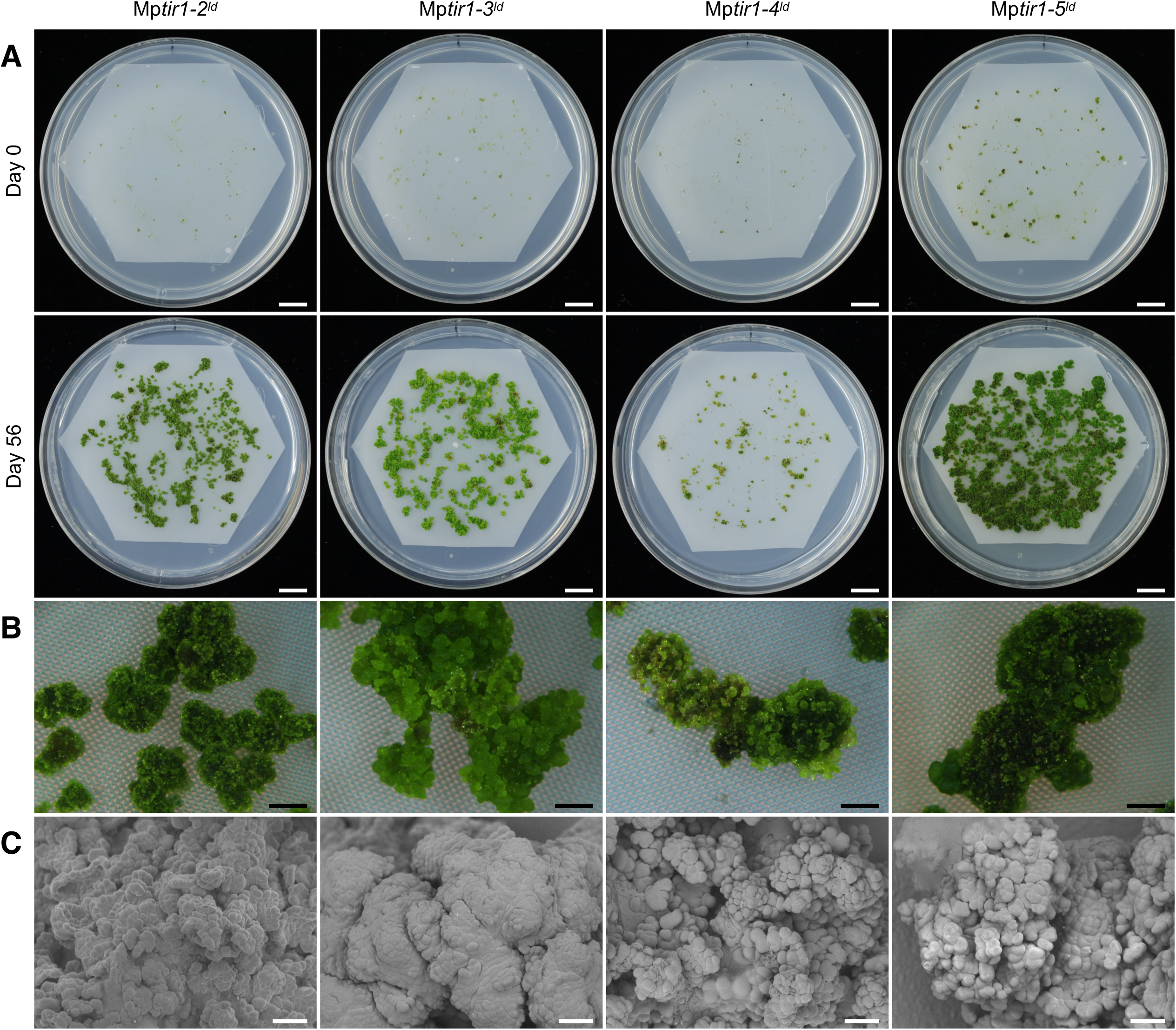
Reproducibility of Mp*tir1* defects in Mp*tir1^ld^* mutants (Supports Figure 4). **(A, B)** Growth of Mp*TIR1*-locus deletion mutants. Mp*tir1^ld^* cells were grown on agar media covered with nylon mesh. Top and bottom panels show identical plates at day 0 and 56, respectively. (B) Magnified images of the 56-day-old plants in A. **(C)** SEM images of the Mp*tir1^ld^* cell masses grown for 35 days. Scale bars = 10 mm (A), 1 mm (B), 0.1 mm **(C)**.

**Supplemental Figure 8.**
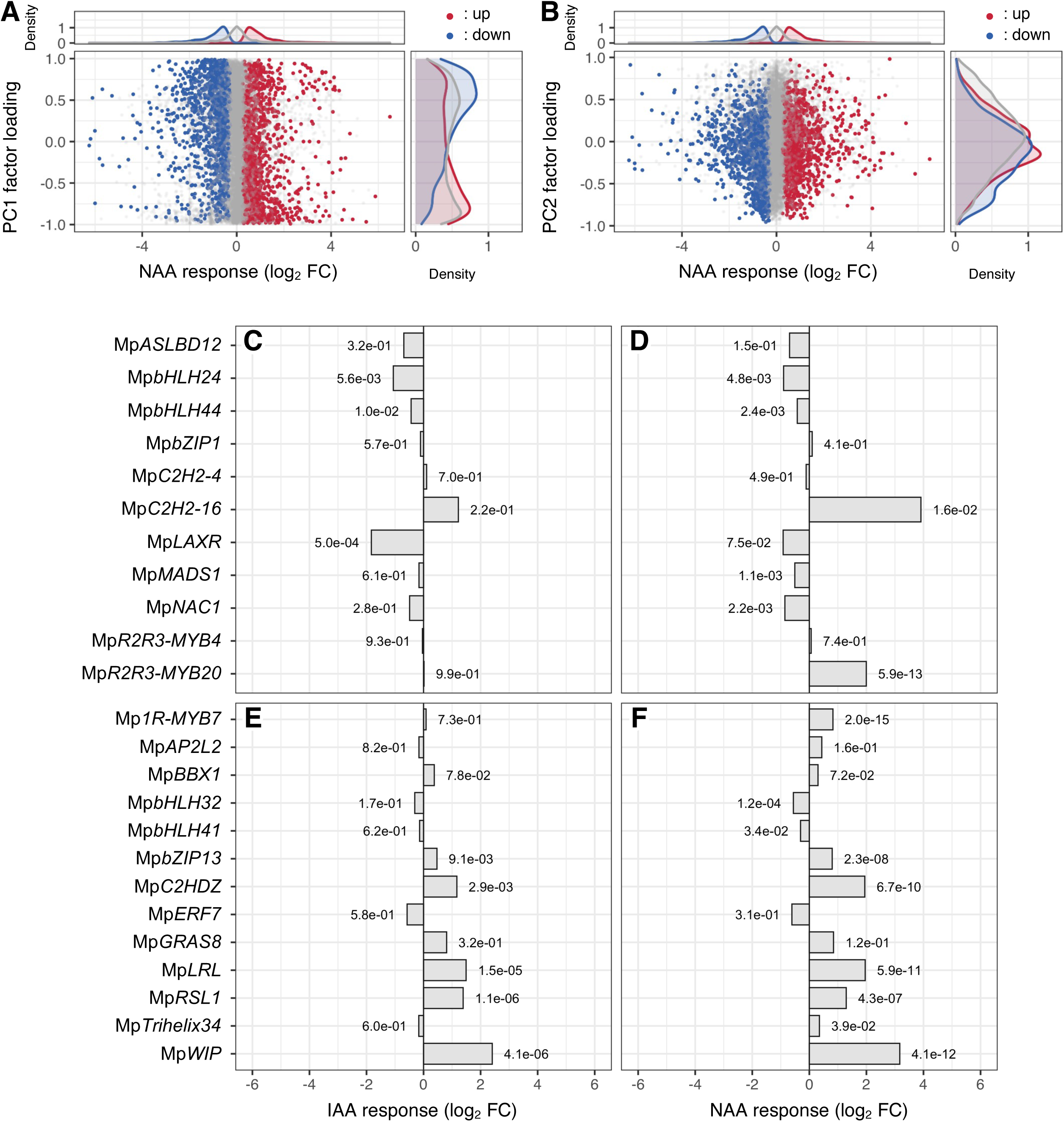
Contribution of auxin-responsive genes to transcriptional properties of Mp*tir1-1^ko^* cells (Supports Figure 7). **(A, B)** Contribution of NAA-responsive genes to factor loadings. X-axis of the central panels represents log_2_ FC upon NAA treatment in WT sporelings. Y-axis of the central panels represents the factor loading of PC1 **(A)** or PC2 **(B)** of the PCA shown in Figure 6B. Red and blue dots indicate significantly (*p_adj_* < 0.001) up- and down-regulated genes upon NAA treatment, and gray dots represent NAA-non-responsive genes. Top and right panels represent distribution densities of the colored dots along x- and y-axes, respectively. **(C–F)** Auxin responsiveness of the differentially expressed transcription factor genes in Mp*tir1-1^ko^* cells (see Figure 6E, F). Transcription factor genes that showed higher **(C, D)** or lower **(E, F)** expression in Mp*tir1-1^ko^* cells in both comparisons with sporelings and thalli are listed. X-axis represents log_2_ FC upon IAA **(C, E)** or NAA **(D, F)** treatment in sporelings. The value beside each bar indicates *p_adj_*.

**Supplemental Table 1.**
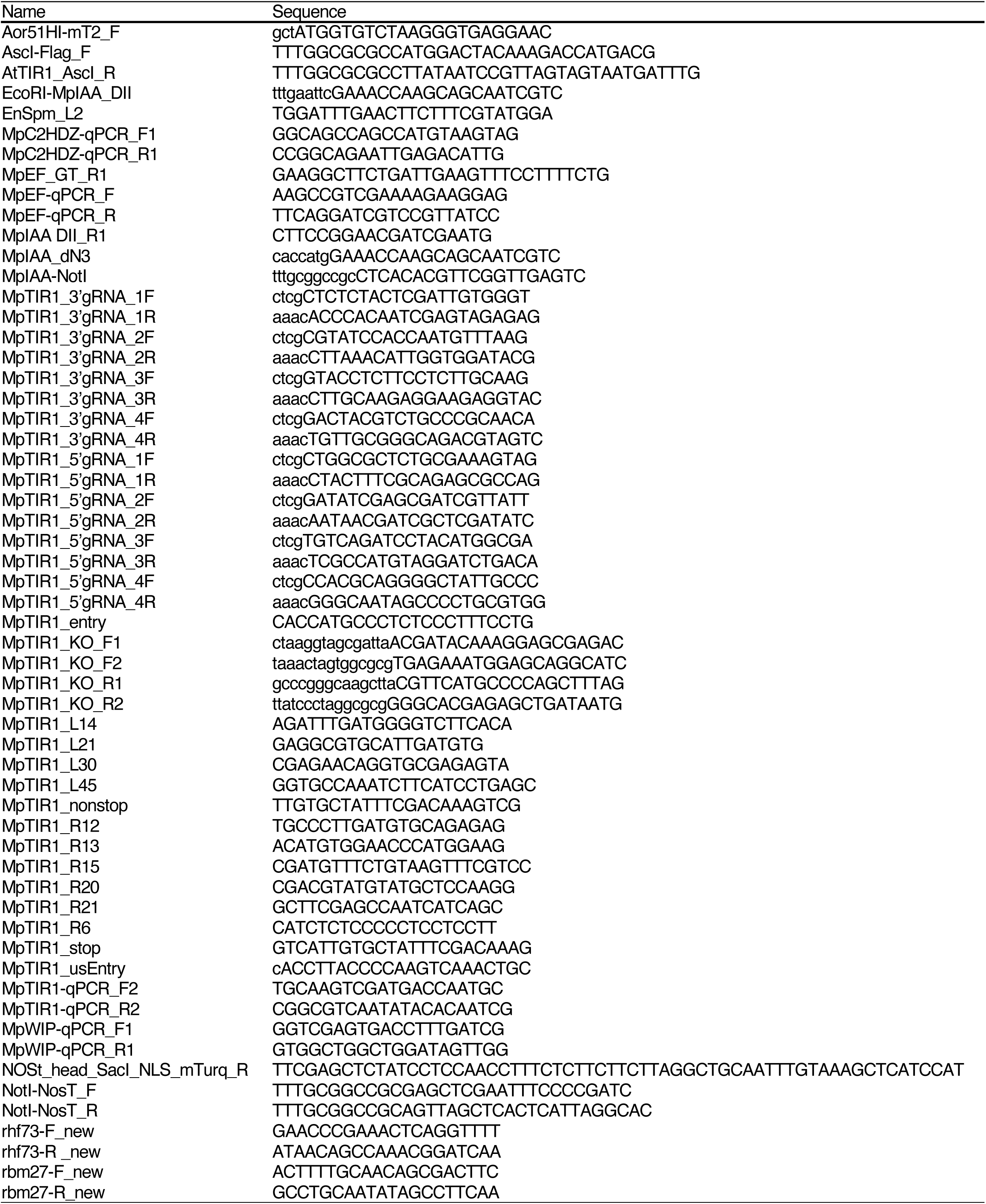
Oligos used in this study.

**Supplemental Table 2.** Public RNA-seq data used in this study.

| ID | Sample | Reference |
| --- | --- | --- |
| DRR050343 | 11-day-old thalli | Higo et al., 2016 |
| DRR050344 | 11-day-old thalli | Higo et al., 2016 |
| DRR050345 | Mature antheridiophores | Higo et al., 2016 |
| DRR050346 | Mature antheridiophores | Higo et al., 2016 |
| DRR050347 | Mature antheridiophores | Higo et al., 2016 |
| DRR050348 | Mature antheridiophores | Higo et al., 2016 |
| DRR050349 | Antheridia | Higo et al., 2016 |
| DRR050350 | Antheridia | Higo et al., 2016 |
| DRR050351 | Mature archegoniophores | Higo et al., 2016 |
| DRR050352 | Mature archegoniophores | Higo et al., 2016 |
| DRR050353 | Mature archegoniophores | Higo et al., 2016 |
| DRR096278 | Gemma cups of 21-day-old thalli | Yasui et al., 2019 |
| DRR096279 | Gemma cups of 21-day-old thalli | Yasui et al., 2019 |
| DRR096280 | Gemma cups of 21-day-old thalli | Yasui et al., 2019 |
| DRR096281 | Midribs of 21-day-old thalli | Yasui et al., 2019 |
| DRR096282 | Midribs of 21-day-old thalli | Yasui et al., 2019 |
| DRR096283 | Midribs of 21-day-old thalli | Yasui et al., 2019 |
| DRR096284 | 7-day-old thalli | Yasui et al., 2019 |
| DRR096285 | 7-day-old thalli | Yasui et al., 2019 |
| DRR096286 | 7-day-old thalli | Yasui et al., 2019 |
| SRR1553294 | Gametophytic apical cells | Frank et al., 2015 |
| SRR1553295 | Gametophytic apical cells | Frank et al., 2015 |
| SRR1553296 | Gametophytic apical cells | Frank et al., 2015 |
| SRR1553297 | Immature sporophytes | Frank et al., 2015 |
| SRR1553298 | Immature sporophytes | Frank et al., 2015 |
| SRR1553299 | Immature sporophytes | Frank et al., 2015 |
| SRR4450254 | 96-hour-old sporelings | Bowman et al., 2017 |
| SRR4450255 | 96-hour-old sporelings | Bowman et al., 2017 |
| SRR4450256 | 96-hour-old sporelings | Bowman et al., 2017 |
| SRR4450257 | 72-hour-old sporelings | Bowman et al., 2017 |
| SRR4450258 | 72-hour-old sporelings | Bowman et al., 2017 |
| SRR4450259 | 24-hour-old sporelings | Bowman et al., 2017 |
| SRR4450260 | 0-hour-old spores | Bowman et al., 2017 |
| SRR4450261 | 0-hour-old spores | Bowman et al., 2017 |
| SRR4450262 | 0-hour-old spores | Bowman et al., 2017 |
| SRR4450263 | 48-hour-old sporelings | Bowman et al., 2017 |
| SRR4450264 | 48-hour-old sporelings | Bowman et al., 2017 |
| SRR4450265 | 24-hour-old sporelings | Bowman et al., 2017 |
| SRR4450266 | 24-hour-old sporelings | Bowman et al., 2017 |
| SRR4450267 | 72-hour-old sporelings | Bowman et al., 2017 |
| SRR4450268 | 48-hour-old sporelings | Bowman et al., 2017 |
| DRR354266 | 5-day-old sporelings (-IAA) | this study |
| DRR354267 | 5-day-old sporelings (-IAA) | this study |
| DRR354268 | 5-day-old sporelings (-IAA) | this study |
| DRR354269 | 5-day-old sporelings (+IAA) | this study |
| DRR354270 | 5-day-old sporelings (+IAA) | this study |
| DRR354271 | 5-day-old sporelings (+IAA) | this study |
| DRR354272 | *Mptir1-1<sup>ko</sup>* (-IAA) | this study |
| DRR354273 | *Mptir1-1<sup>ko</sup>* (-IAA) | this study |
| DRR354274 | *Mptir1-1<sup>ko</sup>* (-IAA) | this study |
| DRR354275 | *Mptir1-1<sup>ko</sup>* (-IAA) | this study |
| DRR354276 | *Mptir1-1<sup>ko</sup>* (-IAA) | this study |
| DRR354277 | *Mptir1-1<sup>ko</sup>* (-IAA) | this study |
| DRR354254 | 5-day-old sporelings (-NAA) | this study |
| DRR354255 | 5-day-old sporelings (-NAA) | this study |
| DRR354256 | 5-day-old sporelings (-NAA) | this study |
| DRR354257 | 5-day-old sporelings (+NAA) | this study |
| DRR354258 | 5-day-old sporelings (+NAA) | this study |
| DRR354259 | 5-day-old sporelings (+NAA) | this study |
| DRR354260 | *Mptir1-1<sup>ko</sup>* (-NAA) | this study |
| DRR354261 | *Mptir1-1<sup>ko</sup>* (-NAA) | this study |
| DRR354262 | *Mptir1-1<sup>ko</sup>* (-NAA) | this study |
| DRR354263 | *Mptir1-1<sup>ko</sup>* (+NAA) | this study |
| DRR354264 | *Mptir1-1<sup>ko</sup>* (+NAA) | this study |
| DRR354265 | *Mptir1-1<sup>ko</sup>* (+NAA) | this study |

